# Local and Global Patterns Support Medical Imaging as a Biomarker of Ageing

**DOI:** 10.1101/2025.05.18.654705

**Authors:** Tamara T. Mueller, Sophie Starck, Rozafe Llalloshi, Georgios Kaissis, Alexander Ziller, Robert Graf, Christopher Schlett, Steffen Ringhof, Fabian Bamberg, Mark Wielpütz, Henry Völzke, Michael Leitzmann, Thoralf Niendorf, Thomas Keil, Lilian Krist, Tobias Pischon, André Karch, Klaus Berger, Jan Kirschke, Daniel Rueckert, Rickmer Braren

**Affiliations:** Lab for AI in Medicine and Healthcare, Technical University of Munich, Germany; Department of Diagnostic and Interventional Neuroradiology, School of Medicine, TUM University Hospital, Germany; Department of Diagnostic and Interventional Radiology, Medical Center, University of Freiburg, Faculty of Medicine, Germany; Institut für Community Medicine, University Medicine Greifswald, Germany; Institute for Epidemiology and Preventive Medicine, University of Regensburg, Germany; Max Delbrück Center for Molecular Medicine, Helmholtz Association, Germany; Institute of Social Medicine, Epidemiology and Health Economics, Charité – Universitätsmedizin Berlin, Germany; Institute of Clinical Epidemiology and Biometry, University of Würzburg, Germany; State Institute of Health I, Bavarian Health and Food Safety Authority, Erlangen, Germany; Institute of Epidemiology and Social Medicine, University of Münster, Germany; Department of Computing, Imperial College London, United Kingdom; Institute for diagnostic and interventional radiology, Klinikum rechts der Isar, Technical University of Munich, Germany; German Cancer Consortium (DKTK), Munich partner site, Heidelberg, Germany; Department of Diagnostic and Interventional Radiology and Nuclear Medicine, University Medical Center Hamburg-Eppendorf, Germany

**Keywords:** Medical Imaging, Ageing Biomarkers, Accelerated Ageing, Population Studies

## Abstract

**Background:** Understanding human ageing across multiple organs is essential for characterising individual health trajectories and identifying abnormal ageing processes. Multiorgan imaging provides an opportunity to quantify biological ageing beyond chronological age. The aim of this study is to assess organ-specific and whole-body ageing patterns and their associations with disease and lifestyle factors.

**Methods:** In this large-scale study, we evaluate biological ageing patterns using 70,000 MRI scans from the UK Biobank and the German National Cohort. We employ 3D ResNet-18 models to predict chronological age from various body regions (brain, heart, liver, spine, lungs, muscle, and intestine) and the whole body. From these predictions, we derive “age gaps” relative to a strictly healthy reference cohort, which enables the identification of accelerated ageing patterns. We then evaluate associations with chronic diseases and lifestyle factors, and a virtual ageing framework was developed to explore counterfactual scenarios by substituting anatomical regions across subjects, quantifying local impacts on global biological age.

**Results:** Here we show significant associations between detected accelerated ageing and specific chronic diseases, including multiple sclerosis and chronic obstructive pulmonary disease, as well as lifestyle factors such as smoking and physical activity. Virtual substitution of anatomical regions demonstrates that local substitutions can influence global ageing patterns.

**Conclusions:** This study demonstrates that multi-organ imaging enables the detection of abnormal ageing patterns at both local and global levels. The presented framework provides a foundation for improved risk stratification and supports the development of personalised approaches to health assessment and disease prevention.

**Plain Language Summary:** As people age, different organs in the body might not always age at the same pace. Understanding these differences can help to explain a person’s health and why they develop diseases earlier than others. In this study, we measure how ageing varies across the body using medical images. We analysed about 70,000 whole-body scans from large population studies in the United Kingdom and Germany. Using AI models, we estimated a person’s biological age from images of different organs and compared it with their actual age. We found that faster ageing in specific organs is linked to certain diseases (such as multiple sclerosis) and lifestyle factors (like smoking and physical activity). These findings may help improve early disease detection and support more personalised approaches to health and ageing in the future.

## 1 Introduction

Ageing is an inevitable aspect of life and the most common risk factor for several diseases [1, 2]. It is a complex interplay between the passing of time, genetic factors, lifestyle characteristics, and environmental impacts [3]. It comes with a decline in functional capacity and stress resilience and affects most or all tissues in the body [4]. One goal of ageing research is to distinguish between the *chronological age*, which refers to the time that has passed since birth, and the *biological age*, aims to describe the actual degree of ageing occuring across cells, tissues, and organs. The aim of estimating a person’s biological age is to obtain a more accurate estimator for risk factors for diseases and mortality [5, 6]. The difference between biological age (extracted from the ageing biomarkers) and chronological age is often referred to as the *age gap*. With this distinction, one can analyse accelerated (biological age > chronological age) or decelerated (biological age < chronological age) ageing (see Figure 1). Accelerated ageing can indicate that a person’s health, environment or lifestyle negatively impact the natural ageing process and has led to more advanced age-related changes in their body compared to peers. However, it is challenging to identify reliable biomarkers for ageing that indeed represent the biological age of a person as there is no clinical reference for this. Therefore, multiple studies have investigated ageing biomarkers in different contexts, including cellular, molecular, and physiological biomarkers [7].

**Fig. 1.**
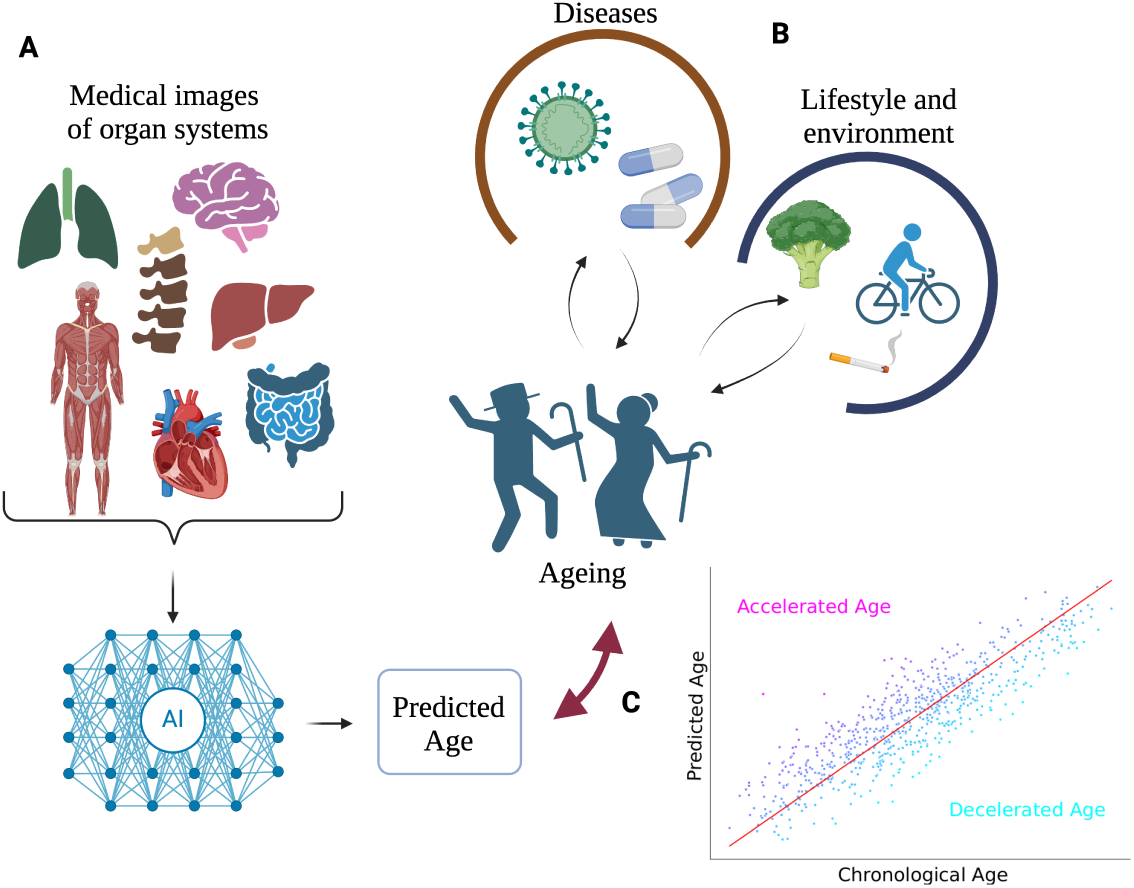
Overview of the whole ageing analysis pipeline. **A:** We investigate how medical imaging can be used to predict ageing of the whole body or individual organ systems and **B:** correlate resulting age gaps with diseases and lifestyle features. **C:** We refer to accelerated age if the predicted age is larger than the chronological age and decelerated age if it is the other way around.

Previous works have defined criteria that a suitable ageing biomarker should meet: Butler et al.[8], for instance, define that ageing biomarkers should be able to predict the outcome of age-sensitive tests better than chronological age and predict longevity at an age at which 90% of the population is still alive. A variety of such ageing biomarkers have been studied and shown promising results in detecting biological age: Burkle et al. [4] introduce MARK-AGE, a project that aimed at identifying a set of ageing biomarkers that measure biological age better than the individual components. They collect physiological data, genomic and protein data, immunological markers, hormones, and metabolic data. Some other works investigate age prediction for individual organs, such as the kidney [9], the lungs [10], or the heart [11], showing promising directions of analysing ageing in more local regions of the human body. Tian et al. [12] were the first to develop a whole-body multi-organ characterisation of ageing. The authors utilise both, non-imaging and image-derived phenotypes with support vector machines.

However, end-to-end deep learning approaches with medical imaging have been used less frequently for age prediction and especially whole-body images are under-explored in this domain. Brain imaging has been, for instance, used to predict brain age and several works have shown how brain age predictions correlate with mortality and (brain-related) diseases [13–17]. Ecker et al. [18] are one of few works using imaging data of different organ systems and deep learning techniques to assess biological age and have shown promising first results in this direction. The authors use different organ systems compared to our study and do not investigate the utility of their age predictions as biomarkers for ageing and the interplay of the different body systems with whole-body ageing.

In this work, we investigate how medical imaging can be used directly as ageing biomarkers and employ end-to-end deep learning techniques to quantify accelerated ageing. We study both local and global components, their interplay, and their connection to lifestyle factors, diseases, and mortality. Furthermore, we introduce a counterfactual “Virtual Ageing Model” framework to simulate individual organ-specific interventions and potentially guide personalised recommendations and interventions. With guided interventions, a slower age-related decline can be targeted, which can lead to more healthy longevity [19].

## 2 Methods

We here summarise the methods utilised in this work, including details about the dataset, the deep learning methods, and the various analyses: such as a organ ageing interplay, lifestyle correlations and Virtual Ageing Model experiments. Furthermore, we provide more details of the selected diseases and non-imaging phenotypes. The deep learning workflow is described in Figure 2.

**Fig. 2.**
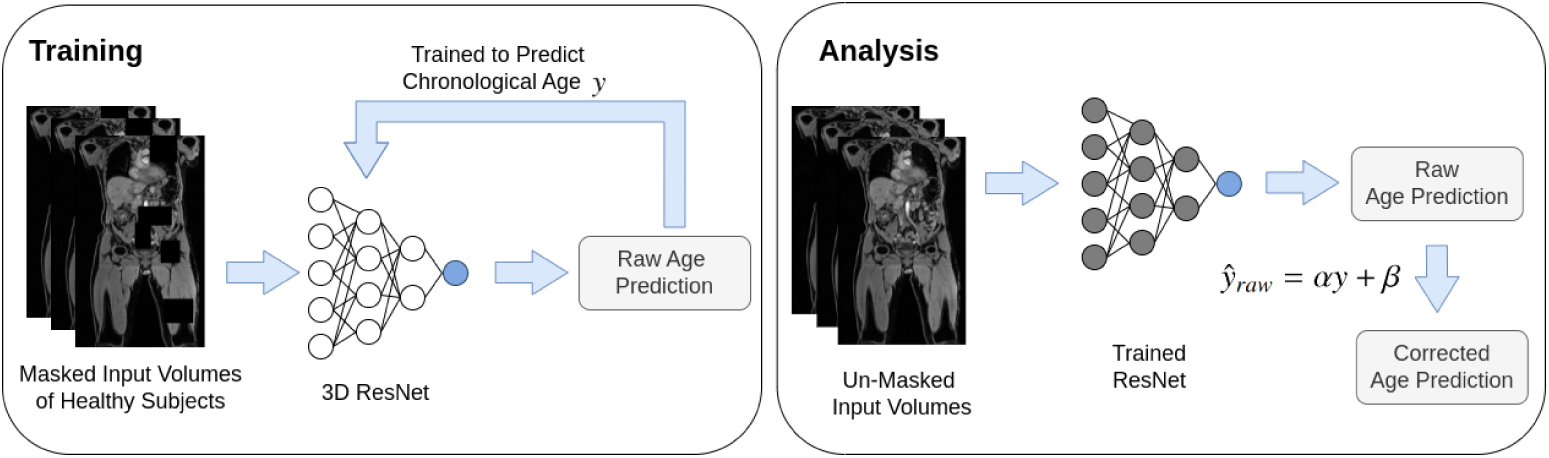
Overview of the biological age prediction model. We train a 3D ResNet18 model for each imaging dataset to predict the chronological age of the healthy training subjects. Here, we use the masked input images. Subsequently, using these raw age predictions, we fit a linear regression model to correct the regression-toward-the-mean bias and obtain the corrected age gap for all subjects. We speak of accelerated ageing, if the age gap is positive and decelerated ageing if it is negative

### Dataset

#### UK Biobank

We utilise (a) 73 094 T1-weighted, dual-echo gradient whole-body MRI datasets with a size of 224 × 168 × 363 voxels and a spatial resolution of 2, 23 × 3 × 2, 23mm. The whole-body MR images were acquired in several stations and stitched using the pipeline of [20]. Furthermore, we use (b) 45 058 T1-weighted scull-stripped brain MRI datasets of the UK Biobank with an isotropic spacing of 1mm^3^ and a size of 160 × 225 × 160 voxels. Although the UK Biobank dataset contains repeat scans for the same participants, we retain only the first one to avoid any data leakage.

We split the dataset into healthy and unhealthy subjects. Healthy subjects are selected as ones that do not have ICD-10 entries or self-reported diseases. “Healthy” subjects are used exclusively for training the model and are divided into a fixed training (80%) and validation (20%) split. The validation set is used solely for model selection and early stopping. “Unhealthy” subjects are held out entirely and used only as a test set to evaluate performance on pathological cases. To assess generalisation, we further introduce an external validation cohort from the NAKO study [21], which is completely unseen during training and model selection. The detailed numbers of samples for all splits and datasets can be found in Table 2. The sample sizes for all training sets are comparable and yield a substantial improvement over the naive mean-age baseline, indicating adequate generalisation. All experiments are repeated five times using different random seeds while keeping the data splits fixed, and results are reported as mean ± standard deviation across runs. We aim for the model to learn to predict the chronological age of healthy subjects and, therefore, be able to identify visual changes in the “unhealthy” subjects of the test set.

In order to investigate images of specific body regions as ageing biomarkers, we use the segmentation pipeline of [22] and select six body regions of interest: the heart, the liver, the spine, the muscle, the lungs, and the intestine. We segment all training and validation samples and 33 135 test samples. We filter out subjects for which the segmentation did not yield any results for specific body regions and exclude these subjects from the pipeline. Furthermore, we dilate the original segmentation mask and select the largest connected component only in order to exclude any faulty segmentations. We then crop the image to the region of interest, containing the region of interest, and mask out all voxels that do not represent the region of interest plus the dilation. The concrete region-wise labels we used are listed in Table 1, column “*Segmentation*”.

**Table 1.**
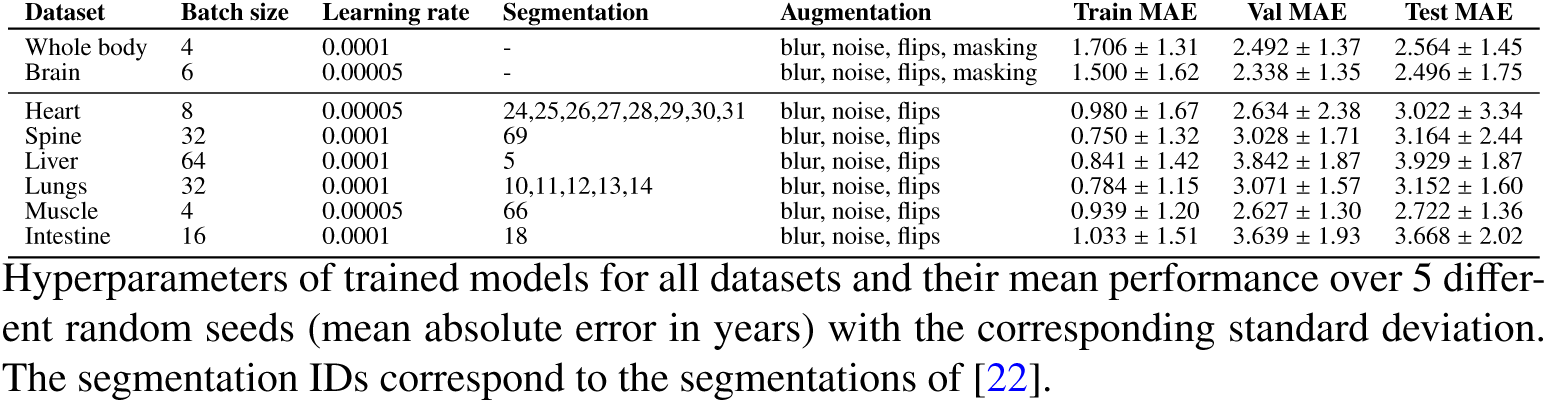
Overview of all hyperparameters and performances for all models.

| Dataset | Batch size | Learning rate | Segmentation | Augmentation | Train MAE | Val MAE | Test MAE |
| --- | --- | --- | --- | --- | --- | --- | --- |
| Whole body | 4 | 0.0001 | - | blur, noise, flips, masking | $1.706 \pm 1.31$ | $2.492 \pm 1.37$ | $2.564 \pm 1.45$ |
| Brain | 6 | 0.00005 | - | blur, noise, flips, masking | $1.500 \pm 1.62$ | $2.338 \pm 1.35$ | $2.496 \pm 1.75$ |
| Heart | 8 | 0.00005 | 24,25,26,27,28,29,30,31 | blur, noise, flips | $0.980 \pm 1.67$ | $2.634 \pm 2.38$ | $3.022 \pm 3.34$ |
| Spine | 32 | 0.0001 | 69 | blur, noise, flips | $0.750 \pm 1.32$ | $3.028 \pm 1.71$ | $3.164 \pm 2.44$ |
| Liver | 64 | 0.0001 | 5 | blur, noise, flips | $0.841 \pm 1.42$ | $3.842 \pm 1.87$ | $3.929 \pm 1.87$ |
| Lungs | 32 | 0.0001 | 10,11,12,13,14 | blur, noise, flips | $0.784 \pm 1.15$ | $3.071 \pm 1.57$ | $3.152 \pm 1.60$ |
| Muscle | 4 | 0.00005 | 66 | blur, noise, flips | $0.939 \pm 1.20$ | $2.627 \pm 1.30$ | $2.722 \pm 1.36$ |
| Intestine | 16 | 0.0001 | 18 | blur, noise, flips | $1.033 \pm 1.51$ | $3.639 \pm 1.93$ | $3.668 \pm 2.02$ |

Hyperparameters of trained models for all datasets and their mean performance over 5 different random seeds (mean absolute error in years) with the corresponding standard deviation. The segmentation IDs correspond to the segmentations of [22].

The UK Biobank has ethical approval from the North West Multi-centre Research Ethics Committee to handle human participant data; no additional ethical approval was required because the study involved the secondary use of data. Written informed consent was obtained from all participants, and all data are deidentified for analysis. Eligible researchers can access UK Biobank data on www.ukbiobank.ac.uk upon registration. For this study, permission to access and analyse the UK Biobank data was approved under the application 87802.

### German National Cohort (NAKO)

For external validation, we use torso MRI data from the German National Cohort (NAKO) [21]. The NAKO images comprise stitched gradient-echo sequences covering the field of view from neck to knee. For comparability to the training dataset, all images were resampled to the same spatial resolution of 2, 23 × 3 × 2, 23mm and size of 160 × 225 × 160 voxels. Anatomical segmentations were also obtained using [22]. The original age distribution of the NAKO spans over 19-74 years, we retain the same range as the UK Biobank 46-83 years for comparability. In this work, the NAKO cohort is used exclusively for external validation and remains completely unseen during model training and internal validation. The German National Cohort was conducted in accordance with national laws and the ethical principles of the Declaration of Helsinki (1975). The study protocol and all human examinations were approved by the Bavarian Medical Association (Bayerische Landesärztekammer) as the central ethics committee. In addition, all participating study centres received approval from their respective local medical associations.

### Deep Learning Models

We train separate models on (1) the neck-to-knee (whole-body) magnetic resonance images (MRIs) of the UKBB, (2) the T1-weighted skull-stripped brain MRIs, and (3) different parts of the whole-body MR images to investigate the ageing of local body regions, using the choronological age as a target. We use 3D ResNet18 for all models due to their established use in related work [15, 23] and their favourable balance between performance and computational efficiency. A consistent architecture also improves the flexibility and generalisability of our approach without requiring additional model selection. For all models, we use the pre-trained 2D weights from ImageNet that are provided by PyTorch [24] and transfer them to the 3D setting by linear inflation along the depth dimension [25]. Several works [26, 27] have shown that one can predict a person’s age successfully from whole-body images. The authors also show that the model specifically focuses on three main regions of interest in the human body: the spine, the aortic region, and the back muscles. In order to prevent the model from focusing on only highly predictive regions in the body, such as the spine and the cardiac area, we randomly mask out parts of the images during training to force the model to use the whole available information to predict a subject’s age. We use this “masking” as a regularisation technique [28, 29] for the whole-body and the brain age predictors, an empirical justification for this training strategy can be found in the Supplementary Material, Figure A1. Furthermore, we use random data augmentation (addition of noise, rotation) applied exclusively to the training set during training for all models to prevent over-fitting. For the region-specific deep learning models, we do not apply any masking, since the areas of interest are already limited to small parts of the whole image and we expect the field of view to be large enough to prevent the model from focusing on highly specific sub-regions. All code is implemented using PyTorch [24] and PyTorch Lightning [30].

We always use five models (initialised with five different random seeds) to make the predictions more stable since individual models can disagree on predictions (model multiplicity [31]) and thus construct an ensemble method [32]. The model performances (averaged over five random seeds), the number of data samples, and the age distributions are summarised in Table 2.

**Table 2.** Description of all datasets.

| Dataset | Age Distribution | Train | Val. | Test | Mean Pred. MAE | Val MAE | Test MAE |
| --- | --- | --- | --- | --- | --- | --- | --- |
| Whole Body | $66.37 \pm 7.95$ | 2841 | 712 | 69541 | 6.62 | $2.492 \pm 1.37$ | $2.564 \pm 1.45$ |
| Brain | $64.88 \pm 7.70$ | 1908 | 466 | 42684 | 6.44 | $2.338 \pm 1.35$ | $2.496 \pm 1.75$ |
| Heart | $\left\{ 66.24 \pm 7.96 \right\}$ | 2841 | 712 | 33135 | $6.63$ | $2.634 \pm 2.38$ | $3.022 \pm 3.34$ |
| Spine | | | | | | $3.028 \pm 1.71$ | $3.164 \pm 2.44$ |
| Liver | | | | | | $3.842 \pm 1.87$ | $3.929 \pm 1.87$ |
| Lungs | | | | | | $3.071 \pm 1.57$ | $3.152 \pm 1.60$ |
| Muscle | | | | | | $2.627 \pm 1.30$ | $2.722 \pm 1.36$ |
| Intestine | | | | | | $3.639 \pm 1.93$ | $3.668 \pm 2.02$ |

Overview of the age distribution of the organ-specific dataset, the number of subjects in each dataset (train, test, and val), the baseline mean absolute error (MAE) of a naive mean age predictor over the whole dataset ("Mean Pred. MAE"), and the corresponding MAE in years of our models on the validation and test sets, averaged over five random seeds. The six organ region datasets contain identical subject, and the corresponding values are consistent across all regions.

### Bias Correction

The age distribution of the utilised dataset is slightly unbalanced, with more subjects between 60 and 70 and fewer younger and older subjects. Our models show a regression-toward-the-mean bias [33, 34], underestimating the older subjects’ age and overestimating the younger ones’ age. We, therefore, perform a bias correction on the predicted ages by fitting a linear regression model to predict the raw age gap *ag_raw_* from the DL model from the chronological age *y* of the subject:

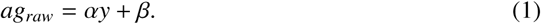

The corrected age gap 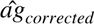 is then defined as the original model prediction *y*^ minus the chronological age *y* minus the fitted age gap:

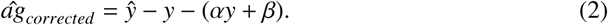

A summary of performances pre- and post-correction can be found in Table A1 of the Appendix. All reported results in this work are based on the corrected age gaps.

### Prediction of Whole-body Age from Body Region Ages

We investigate the interplay between individual body region ages and the whole-body age by fitting (a) a linear regression and (b) a random forest on the region-specific age predictions to estimate the whole-body age prediction. We utilise the same training and test sets as for the deep learning models and use Sklearn’s [35] linear regression and random forest models. The random forest uses a maximum number of 100 estimators and the Poisson criterion. Reported results are coefficient of determination *R*^2^ scores for both the linear regression and the random forest. The *R*^2^ score is defined as 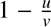, where *u* is the residual sum of squares 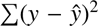 and *v* is the total sum of squares 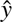), where *y* is the true target value, 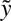 the predicted value and 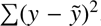 the mean over the true target values. The importance values of the individual input ages (heart age, brain age, spine age, liver age, lung age, muscle age, intestine age) of the linear regression are the coefficients of the fit model. The importance values of the individual input ages (heart age, brain age, spine age, liver age, lung age, muscle age, intestine age) of the random forest are the impurity-based feature importance values.

### Chronic Diseases

We use the UK Biobank fields 41270 for the ICD-10 records and 20002 for the self-reported non-cancer illness codes to determine, whether a subject is counted to be in a specific disease sub-group. We furthermore use the UK Biobank field 41280 to determine the date of diagnosis of the ICD-10 code. We only select subjects that have a record of the specific disease before the time of the imaging assessment or within one year after. Our goal is to evaluate ageing profiles for different diseases, not predicting disease development. Therefore, we are primarily interested in subjects that already show signs of named diseases in their MR images. To account for sample size differences and potential confounding effects, we perform a propensity score weighting [36] on the age gaps for each disease. Propensity scores were estimated separately for each disease using logistic regression with age, sex, weight, height and smoking status (UK Biobank field 20116) as covariates. They were then used to compute stabilised inverse probability weights, which were truncated at the 1st and 99th percentiles to limit the influence of extreme weights. Weighted least squares regression models were then fitted with the biological age gap as the outcome and disease status as the predictor. Effect estimates represent adjusted mean differences in biological age gap between diseased and non-diseased participants. We report adjusted coefficients, 95% CIs and Bonferroni-corrected P-values for the 11 disease groups.

### Lifestyle and Environment

We follow the approach of Bai et al. [37] in performing a phenome-wide association study (PheWAS) with selected non-imaging phenotypes. As confounding variables we use sex, age, sex*age, weight, and height and we remove negative values from the phenotypes since they usually indicate irrelevant responses in the UK Biobank dataset. We furthermore discard features with more than 90% missing data, ones that show more than 95% overlap in entries, and features that correlate higher than 0.9999 with any other utilised feature to avoid redundancies. All features are normalised using mean-standard deviation normalisation for continuous values and rank normalisation for categorical features. We use *P* < 0.00625, Bonferroni-corrected for 264 features ×8 images, and identify the significant features. We use the UK Biobank feature 20116 at the time point of the second assessment (first imaging assessment) to determine whether a subject is a smoker, a non-smoker, or a previous smoker. 0 indicates a non-smoker, 1 a previous smoker, and 2 a current smoker.

### External Validation

To further validate the generalisability of the pipeline, we introduce an external validation set from the NAKO dataset. The previously described models (trained on the healthy UK Biobank data) were used to predict ages on this external validation. To account for the slight domain shift introduced by using a previously unseen dataset, we correct the estimated ages using a linear correction step (see Bias Correction Section). A held-out portion of 10% of the dataset was used to calibrate this correction for the whole body (*n* = 835) and the lungs (*n* = 765). The correction was applied to the resulting 90% of the data (*n* = 7522 for the whole body and *n* = 6893 for the lungs), which was subsequently used to assess accelerated ageing in smokers.

### Survival Analysis

We use the UK Biobank death registry to perform the survival analysis. More information about the death registry can be found at: https://biobank.ndph.ox.ac.uk/showcase/showcase/ docs/DeathLinkage.pdf. We filter for male subjects younger than 65 years old and female subjects younger than 70, following the statistics from the Office of National Statistics [38], using the tables for England and the survival rates in the years 2021-2023. Furthermore, we only select subjects for which we have a data entry for their month and year of birth, to allow for an accurate calculation of the survival. With this, for the brain dataset, 227 individuals have a confirmed death date, for the whole-body dataset, 249 individuals have a confirmed death date, and for the body-region datasets, 106 individuals have a confirmed death date. Living individuals were right-censored, where survival duration was calculated as days between the date of assessment, where the images were recorded, and the date of data download (1st April 2024). The Cox proportional hazards models were performed assuming that the mortality HRs in relation to the age gaps of all datasets stay constant over time. We use the Python package lifelines [39] for the analysis and plotting. We utilise the training, validation, and test sets for this analysis.

### Virtual Ageing Model

In order to investigate the impact of individual accelerated body regions on the overall whole-body age prediction, we assess how region-specific changes in the Virtual Ageing Model of a subject affect their whole-body ageing. We here replace highly accelerated regions from medical images with decelerated regions of other subjects. This replacement of the regions is performed in two steps: (1) the images are registered to each other and subsequently (2) the new region is cut out and transferred to the image of the initial subject.

We follow the approach of Starck et al. [40] for whole-body MR image registration and only register subjects of the same sex to one another. For registration, a (a) *reference image* is selected, to which several (b) *moving images* can be registered. We (a) select a random subject for each sex that does not show accelerated whole-body ageing and evaluate their region-specific age gaps. These subjects have an age of 66.1 years (female) and 64.1 (male). We then (b) select 10 subjects of the same sex that show an accelerated age for each six organ (120 in total). Figure 8B summarises the predicted whole-body and region-specific ages of the selected subjects, which are used to replace the original region of interest in the reference subject (in red). In order to ensure alignment of the individual images, we register the whole-body images of the subjects that will function as the new substituted body regions (moving images) to the selected reference image. We use Deepali [41] as the registration software and register in two steps: First, we affinely align the images and then add a deformable registration step to fine-tune the registration. We note that we cannot replace the brain images but only use the body region from the whole-body images. Subsequently, we cut out the region of interest from the respective moving image and replace the corresponding region in the reference image. We can do this for an arbitrary combination of body regions. In Figure 8, we show an example of artificially altering all accelerated body regions (spine, intestine, muscle, lungs) and compare the whole-body age predictions of the resulting prediction of the Virtual Ageing Model with the original whole-body age prediction of the reference subject over 5 seeds. Additional results can be found in the Supplementary, Section A.5.

### Statistics and Reproducibility

All data analysis was performed with Python 3.8.18 and is reproducible using the code linked below in Code availability. All the data used was obtained as described in the Data Availability section. For all statistical tests, a p-value < 0.05 was considered statistically significant.

## 3 Results

We use (1) 73 094 whole-body and (2) 45 058 brain MR images from the UK Biobank [42] to estimate the biological age of (a) the whole body, (b) the brain, and (c) different localised body areas: muscle, spine, liver, lungs, heart, and intestine. We train convolutional neural networks on a “healthy” subset of the population to predict their chronological age (see Methods Section) and train individual models of chronological age for each body part (whole-body, brain, muscle, spine, liver, lungs, heart, intestine). We refer to the bias-corrected (Section Bias Correction) difference between the predicted age and the chronological age as the *age gap* of a subject.

### Age Predictors

For each imaging dataset (anatomical regions), we train five 3D ResNet-18 models using different randomly initialised networks on a “healthy” sub-population (no ICD-10 records and no self-reported diseases) to predict chronological age. All subjects that have a record of a disease (ICD-10 code or self-reported diseases) are used as test subjects. The concrete numbers of subjects for each imaging dataset can be found in Table 2. Different areas of the human body can potentially age differently [43]. We are therefore interested in *global* ageing patterns –in the whole body and the brain– as well as *local* ageing patterns in different body regions. To investigate the latter, we segment [22] six different body regions in the whole-body MR images and use these regions to predict heart, liver, spine, muscle, intestine and lung age. We here use the same training and validation set as for the whole-body dataset, and a subset of the test set of more than 33 000 subjects.

#### Brain Age Predictor

The model trained on 1908 T1-weighted brain MR images (48% female, 52% male, 66.37 ± 7.95 years) achieves a mean absolute error (MAE) of 2.50 years on the test set (subjects with disease records (ICD-10 code or self-reported)) and 2.34 years on the validation set (subjects without disease records). The MAE of a baseline mean age predictor (which always predicts the mean of the dataset) is 6.62.

#### Whole-body Age Predictor

The mean absolute error (MAE) of the whole-body age prediction model (using the entire neck-to-knee images) is 2.49 years on the validation set and 2.56 years on the test set. The MAE of a baseline naive mean age predictor is 6.44 (47% female, 53% male, 64.88 ± 7.70 years).

#### Body Region Age Predictors

The individual body-region models show generally a slightly higher MAE ranging from 2.72 to 3.93 years than the whole-body (2.56) or the brain age (2.50) predictors since they contain highly local information only. The naive mean age prediction baseline has an MAE of 6.63 years. We note that while we constrain the input of these models to only contain the region of interest in the body, it does not portray a detailed representation of the region of interest and also contains a small portion of the surrounding tissue, due to segmentation inaccuracies and post-processing methods. Further results can be found in the Supplementary Material, including a differentiation between model performances on male and female subjects (Supplementary Figures A2, B8).

#### Impact of Bias correction

Bias correction is performed to mitigate the regression-toward-the-mean effect of ageing datasets. To assess the impact of this step, we analysed both the fitted relationship between the raw age gap and chronological age as well as the resulting prediction errors before and after correction. Supplementary Figure A3 visualises the fitted linear relationship used for bias correction and illustrates the effect of the correction on the regression-to-the-mean bias. In several cases, the correction reduces the age-dependent bias visible in the raw age gap, while in other cases the relationship is weak and the correction has little observable effect. The quantitative impact of the correction is summarised in Supplementary Table A1, which reports the mean absolute error (MAE) for all models on the training, validation, and test sets before and after applying the bias correction. These results indicate that the correction successfully mitigates age-dependent bias where present without negatively affecting model performance when such bias is minimal.

### Interplay of Body Region Ages

Our predicted ages indicate that different areas in the body can age differently. In order to investigate the interplay of the ageing of different body regions, we evaluate their correlation across the whole dataset, which is visualised in Figure 3A. We observe a higher correlation between age gaps in the localised body regions (spine, intestine, heart, liver and lungs). We furthermore note the high correlation between muscle and whole-body age gaps. We suspect that this is because muscle tissue covers large parts of the body, resulting in shared features used for the prediction. Additionally, the lung age and the heart age correlate highly. This might be due to the local proximity of the heart and lungs, causing the heart to be partially visible in the lung images. Moreover, heart and lungs are closely interacting systems, where deoxygenated blood flows from the heart to the lungs to get enriched with oxygen and then back to the heart, from where it is distributed in the body. These two organs are therefore in constant exchange and strongly influence each other. We observe that the brain age gaps are less correlated (≤ 0.34) to all age gaps from the whole body images. This indicates that the brain might age differently from the body and a subject can have an accelerated brain age and a decelerated body age or the other way around. This aligns with related works such as Elliott et al. [44], where the authors have shown that accelerated brain ageing is correlated with accelerated body ageing, but also only loosely. Furthermore, we observe this also in real-world settings, where discrepancies between mental and physical abilities can occur.

**Fig. 3.**
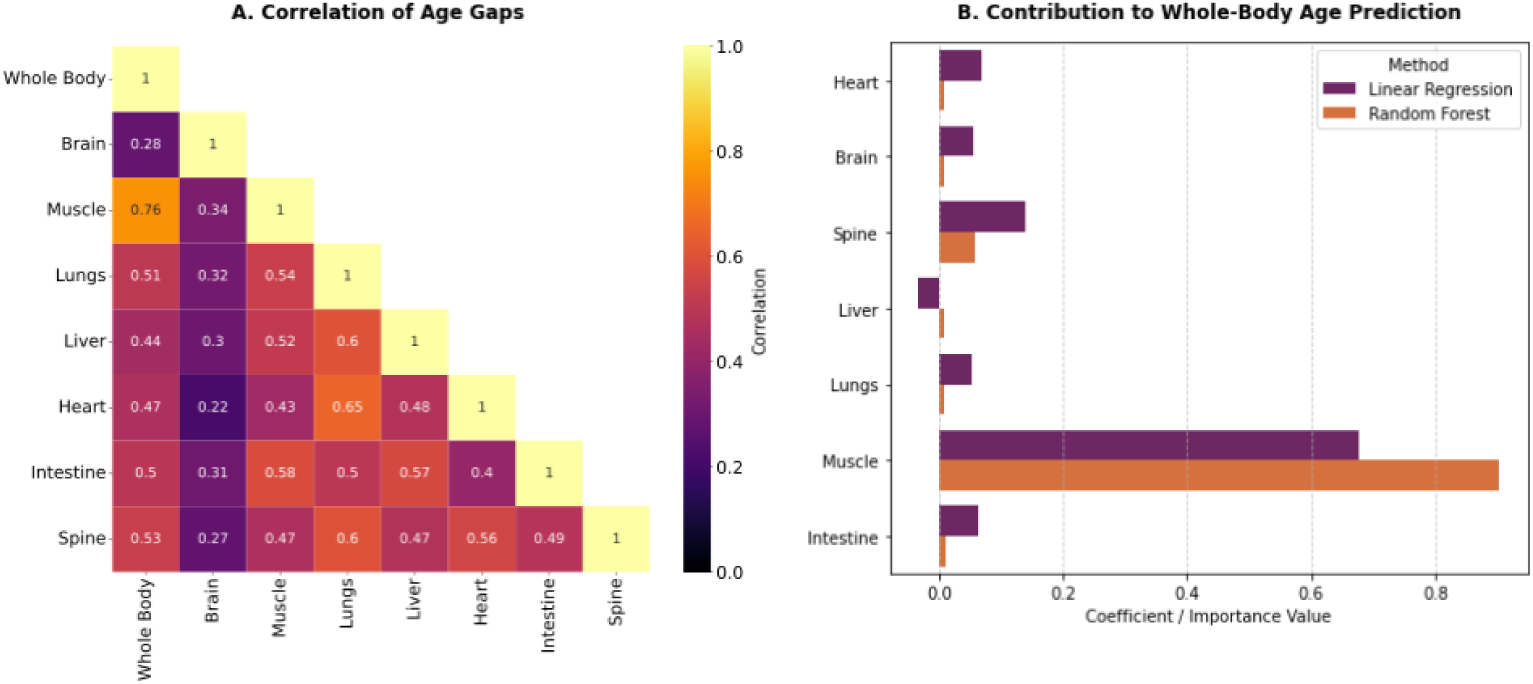
Interplay between regional aging patterns and whole-body age prediction. (A) visualises the correlations between the age gaps of the different body regions. The brain age gaps are less correlated to the body age gaps than the latter among each other, indicating more independent ageing patterns between the brain and the body. (B) quantifies the impurity-based feature importance values of the random forest (*R*^2^ = 0.917) and the coefficients of the linear regression (*R*^2^ = 0.937) for the individual body regions that are used to predict the whole-body age. Aligning with the correlation results, the muscle and the spine ages have the strongest impact on the whole-body age prediction.

Additionally, we investigate how the different body-region age predictions are connected to whole-body ageing. We, therefore, fit a random forest and a linear regression, as very simple but interpretable models. We use all age predictions apart from the whole body (heart, brain, spine, liver, lungs, muscle, intestine) to predict the whole-body age. Subsequently, we evaluate the impurity-based feature importance values of the body-region ages for the random forest and the coefficients for the linear regression to determine the importance of the individual body regions for the whole-body age. The random forest achieves an *R*^2^ score of 0.917 on the test set and the linear regression an *R*^2^ score of 0.937. Muscle and spine ages have the highest importance values and coefficients in predicting the whole-body age from the individual body-region ages (Figure 3B). Interestingly, the linear regression assigns the liver age a slightly negative coefficient. This might be because the liver is a comparably small body region, which might not have a strong impact on the here-evaluated whole-body age or because the liver shows different ageing patterns compared to the whole body. This experiment shows that the prediction of the whole-body age from the individual body-region ages is possible with simple methods and high accuracy. However, we note that ageing is highly complex and that this is only an exemplary way of investigating the interplay between the evaluated region ages in this work on a population level.

### Chronic Diseases

We investigate the interplay between different diseases and accelerated ageing. We select eleven different diseases or disease groups: chronic kidney disease (CKD), multiple sclerosis (MS), diabetes, dementia, stroke, depression, hypertension, ischemic heart disease, scoliosis, chronic obstructive pulmonary disease (COPD), and liver disease (using ICD-10 codes and self-reported diseases from the UK Biobank dataset). For this analysis, we limit the population to be within the 10% and 90% percentile of the age distribution of the training and validation set to reduce the impact of the model bias on the disease-related age gaps. We note that the exact cutoff thresholds are chosen arbitrarily and are only a means to limit the analysis to a range with sufficient data and less model bias. Age prediction models tend to over-estimate younger subjects and under-estimate older subjects [33, 34]. Specific diseases such as dementia and MS are especially prevalent in older or younger subgroups of the population. We furthermore adjust for age, sex, height, weight and smoking using propensity-score weighting [36] to better account for each disease’s effect. More details can be found in Section Chronic Diseases. Figure 4(a) shows the *brain age gap* distributions of named diseases with confidence intervals. Subjects with MS show the highest significant brain age gap of 2.1 years (*n* = 73). This aligns with literature showing that MS patients often show accelerated brain ageing [45, 46]. Other brain-related diseases, such as dementia (*n* = 34) and stroke (*n* = 223) show an increased age gap compared to the validation set, although dementia does not show statistical significance, probably due to the limited sample size. Additionnaly, subjects with body-centred diseases such as scoliosis (*n* = 65), hypertension (*n* = 4994), or heart disease (*n* = 1844) do not show strongly increased brain age gaps.

**Fig. 4.**
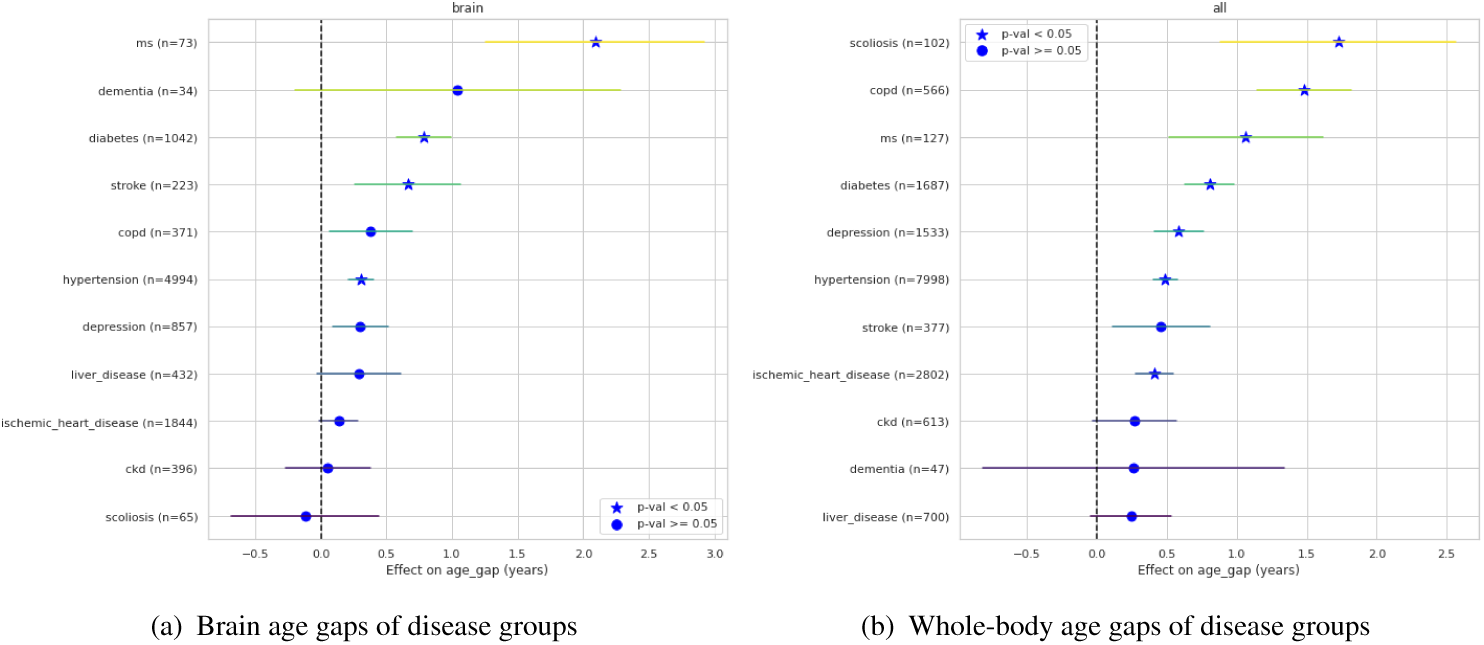
Propensity-weighted age gaps across diseases for brain and whole-body datasets. Propensity-weighted age gaps and CIs for different diseases for (a) the **brain** age predictions and (b) the **whole-body** age predictions. The dotted vertical (black) line indicates no age gap and the stars indicate Bonferroni corrected significant effects (p-value < 0.05). The diseases are ordered by ascending age gap.

Figure 4(b) shows the age gap distributions of the *whole-body age* of the selected diseases. Subjects with scoliosis (*n* = 102) are identified as the sub-group with the highest median whole-body age gap of 1.9 years, followed by COPD subjects (*n* = 566) and MS subjects (*n* = 127). Other works have shown that ML models trained on whole-body images for age prediction have a strong focus on the spine as a region of interest [27]. This might be a factor for scoliosis subjects showing the “strongest” accelerated ageing profile. When analysing disease-specific age gaps for the smaller body regions (lungs, liver, spine, intestine, heart, and muscle), we do not observe a clear correlation between body-region-specific diseases and their age gaps (Section A.2). While subjects with scoliosis show accelerated spine ageing and COPD subjects show accelerated lung ageing, liver-disease subjects do not show accelerated liver ageing and heart-disease and hypertension subjects do not show accelerated heart ageing. There is a multitude of potential reasons for this. For instance, specific diseases might show an impact on the visual appearance of the body region, but this change might not correspond to healthy ageing patterns. Furthermore, the body regions are not independent and while COPD manifests mostly in the lungs, it is often caused by smoking [47], which can affect multiple body regions [48, 49].

### Lifestyle and Environment

We analyse the interplay between different lifestyle and environmental factors (non-imaging) with age gaps of all imaging datasets by performing a phenome-wide association study (PheWAS). We select a set of 264 non-imaging phenotypes of the UK Biobank of 17 different categories: alcohol, body composition, diet, employment, family history, female- and male-specific factors, general health, medical conditions, mental health, pain, physical activity, sexual factors, sleep, smoking, social support, and sun exposure (see Supplementary Material). Figure 5A shows the Manhattan plot of the negative log of the *P*-values (two-sided t-test, *P* < 2.37 × 10^−4^ Bonferroni-corrected for 264 features × 8 image datasets) of the non-imaging phenotypes and the organ-specific predicted age gaps. The PheWAS results show several statistically significant differences between non-imaging phenotypes and the predicted age gaps of different body parts. Some examples are “fat-free muscle mass” and “muscle fat infiltration” from the body composition features, which show significant differences for all investigated ageing types. Several physical activity features show a significant effect for whole-body, lung, liver, muscle, spine, and brain age gaps. Features indicating general health show a significant difference for the age gaps of all body regions. The dietary features that show a significant difference are whether a dietary change has happened within the last five years, cereal intake, and coffee intake. Some social factors (social activities and loneliness) show significant differences for accelerated ageing in almost all datasets. This aligns with findings from studies such as [50].

**Fig. 5.**
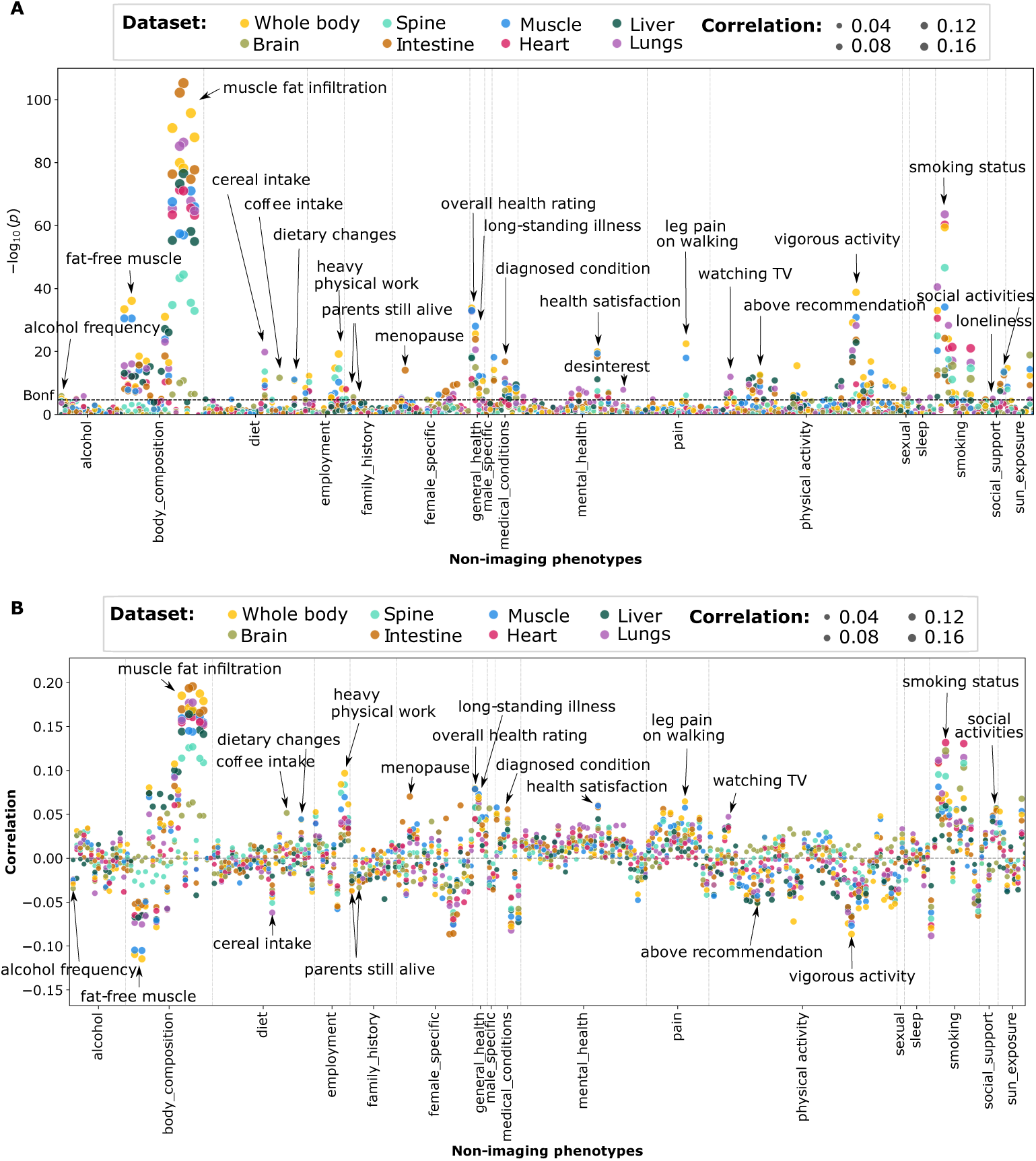
Results of PheWAS study on non-imaging phenotypes and the age gap for different body parts. **A:** Manhattan plot showing the -log10-transformed *P*-values from the PheWAS. The horizontal dashed line indicates the Bonferroni-corrected significance threshold. All association tests are two-sided. **B:** The positive and negative Pearson product-moment correlation coefficients between the non-imaging features and age gaps of the different body parts. Correlations reflect the direction and strength of association, with two-sided significance tests.

The corresponding correlations (positive and negative) are visualised in Figure 5B. We note that a high value in the UK Biobank feature “alcohol frequency” (ID 1558) indicates rare alcohol consumption. The negative correlation therefore indicates that *frequent* alcohol consumption is *positively* correlated with accelerated ageing. A high fat-free muscle mass is negatively correlated with accelerated ageing, while a high muscle fat infiltration is positively correlated with accelerated ageing. We note that possibly some of these non-imaging features are confounders. People who have an overall low score on “health satisfaction”, e.g. might also tend to not be able to exercise a lot or experience more pain.

We identify smoking as one of the most correlated lifestyle factors with higher age gaps (Figure 5B). Figure 6A visualises the age predictions of smokers and non-smokers and their chronological age for the brain, the whole body, and the lungs. Subjects who are current smokers (red) tend to have a higher age gap than non-smokers. This trend is especially visible for the whole body and the lung predictions while being weaker for the brain age prediction. This aligns with the PheWAS analysis, where the brain age gap is less correlated to the smoking status than the age gaps of the other body regions. The mean whole-body age gap of smokers is +1.728 years (*n* = 2254), while the mean age gap of non-smokers is +0.029 years (*n* = 42047). Subjects that have previously smoked, show a mean age gap of +0.380 years (*n* = 23026), supporting clinical findings that cessation of smoking leads to normalisation of e.g. risk for lung cancer [51].

**Fig. 6.**
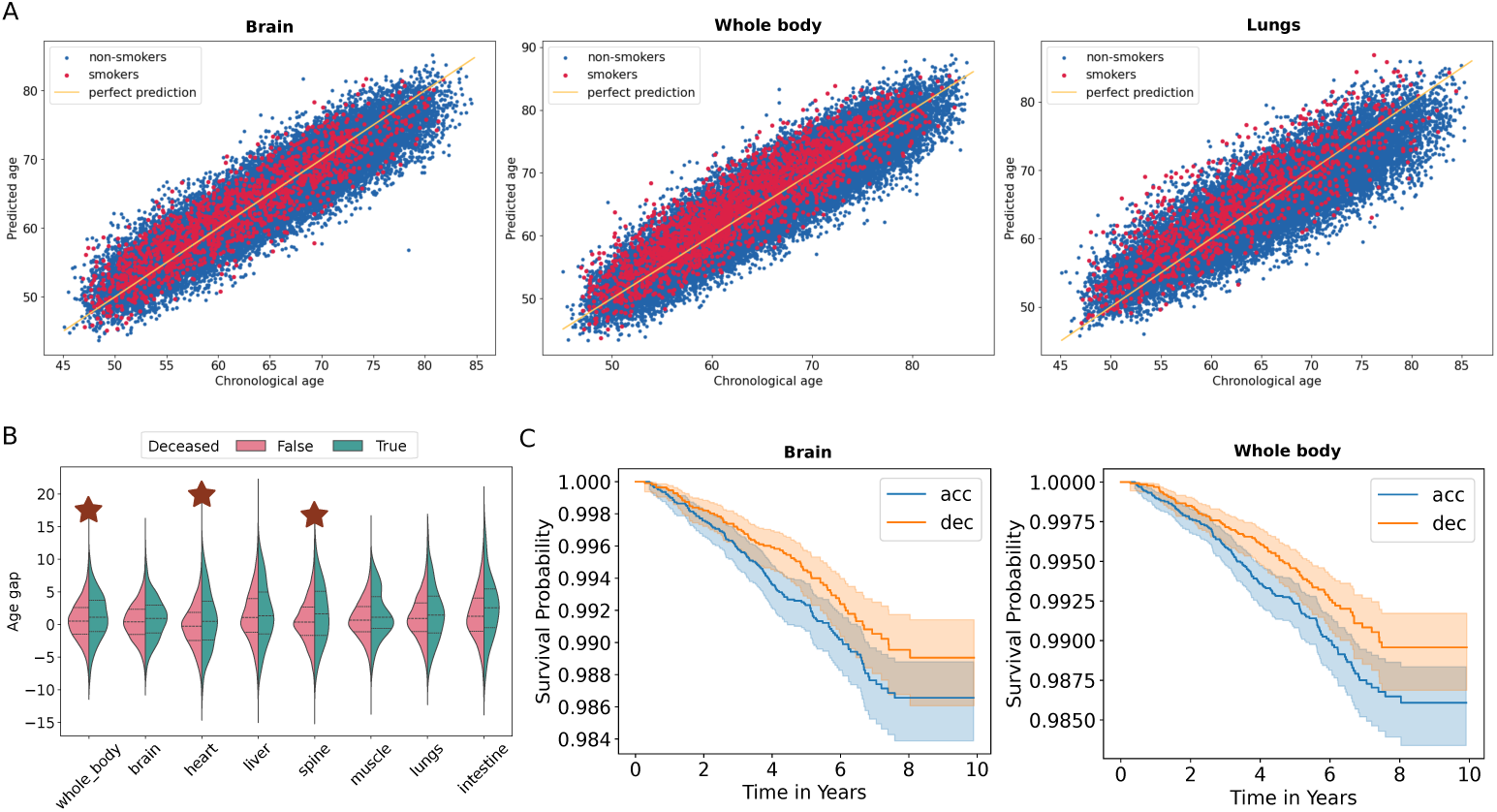
Smoking, age predictions and survival outcomes. **A:** Age predictions (y-axis) vs. chronological age (x-axis) of smokers (red) and non-smokers (blue) for the brain, the whole body, and the lung age (left to right). We see a stronger shift for the whole body and lung age predictions compared to the brain age. **B:** Difference in distributions for subjects which are deceased and ones that are still alive of the population younger than 70 at imaging time point. Statistically significant differences are indicated by the red asterisk (*p* < 0.00625, two-sided t-test, Bonferroni-corrected). Significant comparisons include whole body (*p* = 0.003), heart (*p* = 0.003), and spine (*p* = 0.001). **C:** Kaplan Meier Survival Curve for brain age prediction (left) and the whole-body age prediction (right) for the accelerated agers (blue) and the decelerated agers (orange) with the associated error bands.

### External Validation

To assess the generalisability of the models and analyses and assess the reproducibility of our findings we introduce an external validation on the German National Cohort (NAKO) [21]. The trained models were applied on this independent dataset and a linear correction step was performed to account for systematic prediction biases (details are provided in Section External Validation). Figure 7 shows the predicted ages for smokers and non-smokers for both whole body and lungs ages on NAKO. Results were consistent with the UK Biobank analysis (Section Lifestyle and Environment), with smokers exhibiting higher age gaps than non-smokers for both whole-body and lungs. Whole-body age gaps were +0.557 years in smokers (*n* = 1271) versus −0.215 years in non-smokers (*n* = 6077), while lung age gaps were +0.727 years (*n* = 1177) versus −0.310 years (*n* = 5571), respectively. Linear correction reduced systematic prediction biases, shifting whole-body age gaps from −0.848 to −0.215 years (non-smokers) and +0.824 to +0.557 years (smokers), and lung age gaps from +3.493 to −0.310 years and +5.286 to +0.727 years respectively.

**Fig. 7.**
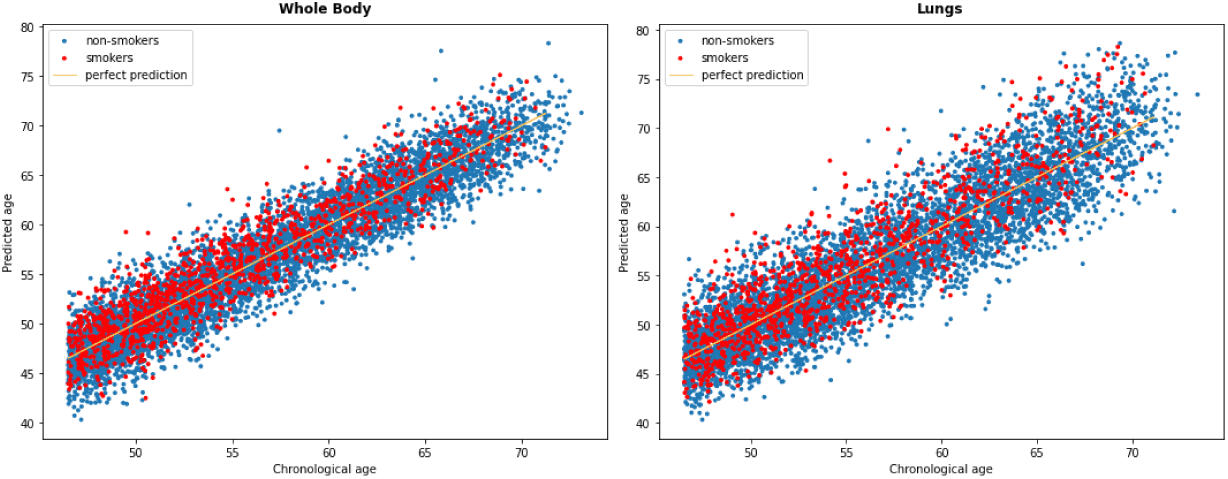
Body and lung age predictions in an external cohort (NAKO) Age predictions (y-axis) vs. chronological age (x-axis) of smokers (red) and non-smokers (blue) for the whole body, and the lung age (left to right) on the NAKO external validation set. We see a similar shift for the whole body and lung age predictions compared to the UK biobank results Figure 6.

### Survival Analysis

To investigate the interplay between the obtained age gaps for different body regions and survival, we utilise data linkages to national death registries in the UK, which are provided by the UK Biobank. Butler et al. [8] define a useful age biomarker as one that predicts longevity at an age at which 90% of the population is still alive. Given the small number of events (*n* = 79 females, *n* = 42 males for the whole body; *n* = 91 females, *n* = 35 males for the brain), we slightly increase this to an age where 85% of the population is still alive for this analysis (*n* = 150 females, *n* = 99 males for the whole body; *n* = 140 females, *n* = 87 males for the brain). Individuals without an entry in the death registry are right-censored (whole body: *n* = 25546 females, *n* = 13739 males; brain: *n* = 17511 females, *n* = 9387 males).

We first compare the age gap distributions between deceased subjects and subjects that are still alive for all body regions. This allows us to study whether the predicted age gaps are correlated to early death. Figure 6B visualises the different age gap distributions for all image datasets. We see an upward shift in the median age gap for the deceased subjects in all image regions and the differences for the whole-body, the heart, and the spine age gaps are significant (indicated by the asterisk) using a two-sided t-test, *P* < 0.00625, Bonferroni-corrected for 8 imaging datasets. While the difference between distributions is non-significant for some image regions (brain, liver, muscle, lungs, intestine), the Cohen’s *d* effect sizes are similar for all body regions apart from the liver (brain: 0.17, whole-body: 0.20, heart: 0.25, liver: 0.07, spine: 0.26, muscle: 0.21, lungs: 0.13, intestine: 0.19).

Secondly, we visualise the Kaplan Meier curves for accelerated (“acc”, blue) and decelerated (“dec”, orange) agers in Figure 6 C for the brain (accelerated age: *n* = 10556, decelerated age: *n* = 7156) and the whole-body age (accelerated age: *n* = 11783, decelerated age: *n* = 8383). For both image datasets, the accelerated agers constantly have a lower survival probability and the shaded confidence interval overlaps less for the whole-body age, aligning with the t-test results from above. Survival durations were right-censored for individuals without an entry in the death registry. Corresponding visualisations for the other image datasets can be found in the Appendix. Finally, we fit a Cox proportional hazard model to the dataset using age, sex, and a binary variable indicating whether a person shows accelerated or decelerated (brain or whole-body) ageing as confounders. The Hazard Ratios (HRs) for the binary attribute indicating accelerated or decelerated ageing is 1.24 for the brain and 1.23 for the whole-body age (*P* < 0.005). Both HRs indicate an increased hazard if the subjects show accelerated ageing.

### Virtual Ageing Model

In order to investigate ageing patterns of different regions in the body for individuals, we construct a “Virtual Ageing Model” – a digitised representation of a person using the MR images of the different body regions and their age predictions. This Virtual Ageing Model (VAM) allows (a) a detailed investigation of existing ageing patterns and (b) artificially changing the appearance of the person and studying the impact of local changes on the whole body on an individual level.

Figure 8A visualises an example workflow of a female subject (a random subject that shows accelerated whole-body age). For a subject that obtains MR images at a medical screening, we can create their VAM. This subject shows an overall whole-body accelerated ageing of +5.9 years. A further investigation of the body regions shows almost *no accelerated ageing* (age gap ≤ 1 year) in the brain, the heart, and the liver, and *mildly accelerated ageing* (age gap ≤ 4 years) in the lungs, the muscles, and the intestine. The most accelerated body-region of this subject is the spine with an age gap of +5.8 years. More results, including artificially changing the appearance of other body regions, are in the Appendix, Section A.5 and Table A2.

**Fig. 8.**
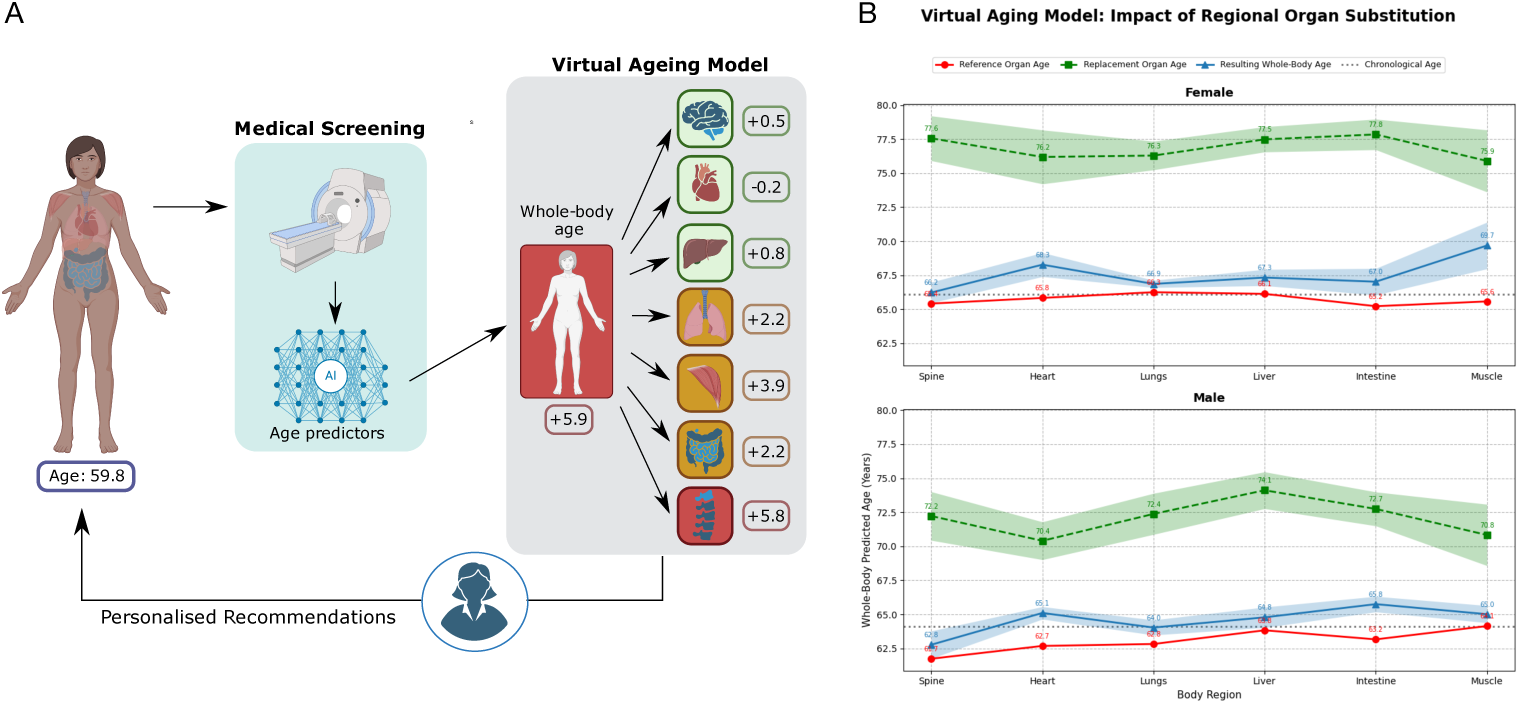
Virtual Ageing Model. (A) is an example screening scenario.A female subject (left) of age 59.8 years obtains MR images at a screening. The Virtual Ageing Model of this subject renders an accelerated whole-body age (+5.9 years) with different accelerations in different body regions. The spine shows the biggest acceleration. This information could be used for personalised recommendations to the patient to consider their region-specific ages in daily life. (B) displays the impact of region-specific and whole-body ages of selected subjects that are used to replace the reference’s body regions. The red line show the reference organ age, in green the age of the new organ substituted and finally in blue the resulting whole body age for each substitution. For each organ 10 accelerated subjects are selected and the whole-body predicted age is evaluated over five seeds.

To evaluate the contribution of individual organs to whole-body predicted age, we applied the VAM as a perturbation-based analysis, replacing each organ independently with an accelerated one. Figure 8B shows the resulting whole-body predicted age (blue), alongside each subject’s reference organ age (red) and the accelerated replacement organ age (green), stratified by sex. For both males and females, accelerated organ substitution consistently increased the whole-body predicted age. In females, heart substitution elevated whole-body age to 68.3 years, while muscle substitution produced the largest shift, reaching 69.7 years. Substitutions of the spine, lungs, liver, and intestine resulted in comparatively modest elevations. In males, a similar pattern emerged: heart, lungs, and intestine substitutions each elevated whole-body predicted age, with muscle again producing a notable upward effect. Across both sexes, however, the absolute increase in whole-body predicted age remained modest relative to the magnitude of the substituted organ age.

These individual ageing patterns could subsequently be used by physicians to formulate informed personalised recommendations. In the PheWAS analysis (Section Lifestyle and Environment), we have, for instance, detected that the frequency of vigorous physical activity is negatively correlated with accelerated spine age. This subject could potentially try to increase their activity and therefore support their spine ageing. On the other hand, some lifestyle factors such as heavy physical work (which is positively correlated with accelerated spine age) might not be adjustable and the subject could be assured that their accelerated whole-body age mostly stems from an accelerated spine age and that the other body regions show more comparable ageing to peers. We note that this is a hypothetical scenario that requires further and more detailed investigations of causality and the impact of specific interventions on the ageing of a subject. We do not perform a de-ageing on the Virtual Ageing Model but investigate the impact of individual body regions on the whole-body age on an individual level, compared to the population-level analysis in Section Interplay of Body Region Ages. As a course of personalised medicine, this analysis would need to be performed for each individual separately and each recommendation needs to be tailored to the subject circumstances, such as lifestyle and ageing patterns. Furthermore, we note that diseases or injuries could also potentially impact the accelerated age. However, such findings could open opportunities for a wide range of interventions, including specific adaptations of working conditions or lifestyle elements.

## 4 Discussion and Conclusion

Several works have previously investigated ageing biomarkers, using various modalities [4, 12, 14, 15]. While brain age has been studied in more detail [13–17], the interplay between brain ageing, whole-body ageing, and the ageing of local regions in the body is underrepresented. Tian et al. [12] show how individual organ systems can shed light on the complex and multi-faceted ageing process in the human body. We show that medical images combined with ML methods can be valid biomarker candidates for ageing. With the full 3D volume of MR images, we achieve accurate age gap predictions and therefore reliable results regarding accelerated and decelerated ageing. Using images eliminates the necessity of selecting features and relying on e.g. self-reported features, both prone to introduce human bias. Moreover, our approach is highly general and can be applied to arbitrary input data, including higher resolution or functional imaging data. We show that brain age is less correlated to the ageing of the whole body and the selected local regions, indicating a more detached ageing process between body and brain. Finally, by creating a Virtual Ageing Model for a subject, we can detect their most accelerated body regions, which can guide recommendations for interventions and lifestyle changes and potentially be incorporated into clinical workflows, as a step towards more personalised medicine.

We note that the age gaps presented in this work are in general smaller than the ones in non-imaging studies such as [12]. This might be because the ML models trained on the images show better age prediction performance and do not have strong deviations from the actual age, even if a disease is present. We opt to interpret the actual number of accelerated years more as a relative measure for “how strongly” someone shows accelerated ageing than for a concrete measure of how many years this person might die earlier. Our models have only seen individuals without disease records during training and have therefore learned to identify healthy ageing patterns. Some diseases might not show the same ageing patterns and therefore might not show more accelerated ageing compared to healthy subjects. Circling back to the defined characteristics of [8] that ageing biomarkers should meet, we can conclude that medical images are good biomarker candidates for ageing but they are no perfect biological age indicators. The discovered correlations between age gaps and lifestyle features highlight several correlated phenotypes. For instance, physical activity, body composition, diet, health status, and social components show significant differences (*P* < 2.37 × 10^−4^, two-sided t-test, Bonferroni corrected for 265 features ×8 images).

While our work renders interesting results about global and local ageing patterns in medical images, we note some limitations. Firstly, the dataset has a limited age range with most subjects centered around 60-70 years old, which makes age prediction especially challenging for under-respresented age groups. This is a very common issue in such ageing datasets that most works address by adding a bias correction step to the pipeline as an effort to mitigate the effects of such a narrow and unbalanced age range [52]. However, an investigation of a dataset with a wider age range would be highly interesting. We furthermore employ a single, uniform network architecture across all organs to ensure comparability between regions. Our goal is to demonstrate the feasibility of a unified framework for analysing region-specific ageing patterns rather than to optimise individual organ predictors. Investigating organ-specific architectures could further improve performance and is an interesting direction for future work.

We also highlight the complexity of the ageing process, with pathological conditions superimposed and resulting from a multitude of impact factors that can change over time. These potential confounding variables are not taken into account in this analysis. In future modelling approaches, other components of the ageing process, such as genetics, the immune system, or behavioural components should be taken into account, to frame a more diverse and robust ageing profile for individuals. Additionally, pathological and physiological ageing phenotypes cannot be distinguished with this method. The age predictors are designed on a “healthy” cohort to model normative ageing patterns, but deviations from this baseline may arise from natural biological variation but also subclinical conditions or undiagnosed diseases. Therefore, an accelerated age should be interpreted as indicating a divergence from healthy ageing trajectories, not as a diagnostic marker or a causal indicator of a disease, which would require future work, including long-term follow-up data and targeted studies. We define our “healthy” cohort as subjects that do not have an ICD-10 record and no self-reported diseases, which might not be a sufficient characteristic to ensure their ageing is “healthy”, as some healthy subjects may harbour undetected conditions. On the contrary, this strict criterion excludes subjects with hypertension, a highly prevalent but highly treatable condition in older subjects, which might not greatly interfere with a healthy ageing process, although its association with an increased risk of stroke and heart disease [53]. Despite its limitations, this is a commonly used approach in current studies [12]. Finally, because our study is cross-sectional, it cannot establish causality or model temporal ageing trajectories, highlighting the need for longitudinal imaging to investigate how ageing patterns evolve within individuals.

Lastly, we note that while imaging techniques allow for a global and general investigation of ageing, obtaining the images is expensive and sometimes difficult to include in standard medical procedures. Still, medical images are used in practice for the assessment of patients on several occasions and show promising results to function as biomarkers for ageing. Region-specific imaging datasets with a higher resolution could be beneficial for such analyses. While we also investigate local ageing patterns by using specific body regions and evaluating their ages, the strong correlation between the age gaps of all body regions shows that with this method, the ageing process cannot be completely separated into different smaller areas in the body. This is expected because most lifestyle factors, as well as diseases, have an impact on not only a single organ. We envision the investigation of more body regions using pipelines such as [22] to get an even better understanding of locally different structures of ageing as interesting future work. We furthermore see the investigation of longitudinal data and the development of ageing within one subject over time as an interesting next step. This could help identify early markers of ageing, inform personalised interventions and monitor the efficacy of such targeted therapies. The routine assessments of the ageing patterns of a single subject, are discussed in form of the here presented Virtual Ageing Model. We note that the VAM does not aim to provide a biological simulation of ageing but merely to explore perturbation-based counterfactual scenarios and to shed light on possible risk factors and potentially guide medical examination towards which regions show especially large age gaps.

## Data Availability

Eligible researchers may access UK Biobank data on www.ukbiobank. ac.uk, upon registration. For this study, permission to access and analyse the UK Biobank data was approved under the application 87802. This project was conducted with data (Application No. NAKO-732) from the German National Cohort (NAKO) (www.nako.de). The NAKO is funded by the Federal Ministry of Education and Research (BMBF) [project funding reference numbers: 01ER1301A/B/C, 01ER1511D, 01ER1801A/B/C/D and 01ER2301A/B/C], federal states of Germany and the Helmholtz Association, the participating universities and the institutes of the Leibniz Association. We thank all participants who took part in the NAKO study and the staff of this research initiative. Figure 1 was created in BioRender. Mueller, T. (2025) https://BioRender.com/8ryioo7. Figure 8A was created in BioRender. Mueller, T. (2025) https://BioRender.com/8rm2rn2

## Code Availability

The source code will be made public upon publication of this work.

## Supplementary Material A Additional Results

### A.1 Model Performance

All model performances are visualised in Supplementary Figure A2. The left column shows the performance on the training set, the middle column on the validation set, and the most-right column on the test set. We note that all models tend to overfit on the training set but still generalise well to the validation and the test set. The degree of overfitting is further characterised in Supplementary Figure A1, which shows the residual distributions for the masked and unmasked whole-body age models across training and validation splits. The masked model exhibits closer agreement between splits, supporting its use as a regularisation strategy. The training and validation sets contain subjects without any records of ICD-10 codes and self-reported diseases and are therefore meant to represent healthy ageing patterns. Additionally, the results pre- and post-bias correction, are reported in Table A1 and visualised in Supplementary Figure A3.

**Supplementary Figure A1.**
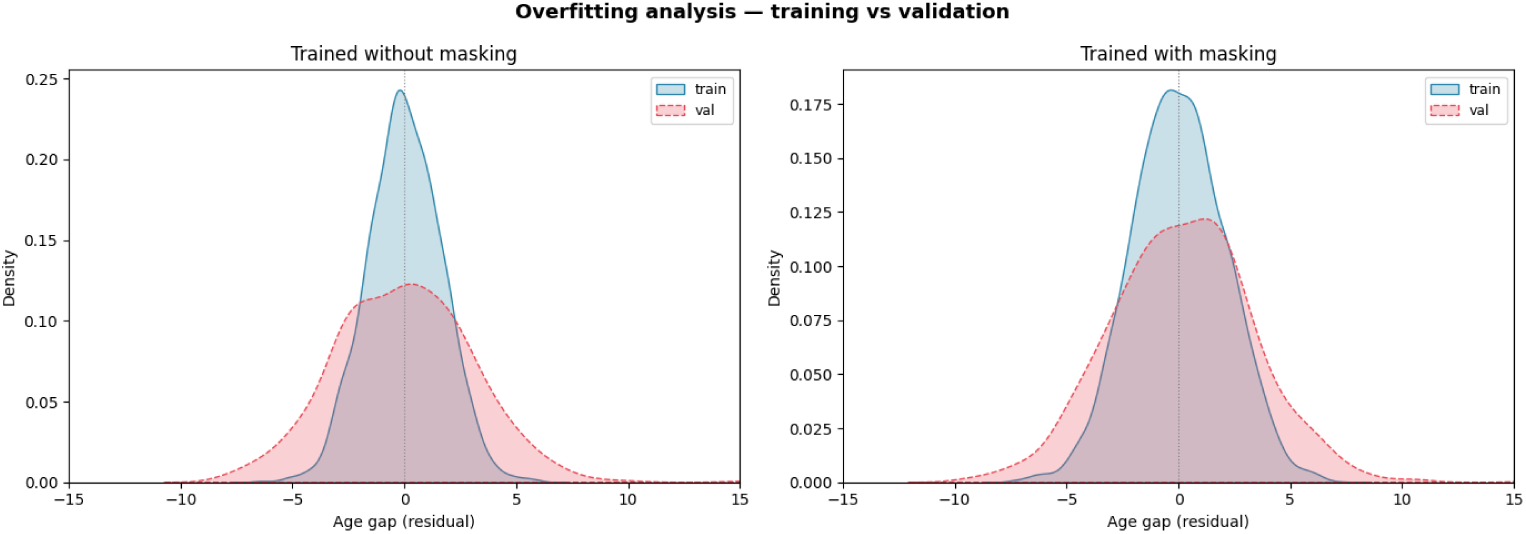
Residual distributions for the masked and unmasked whole-body age models. Kernel density estimates of the residual age gap are shown separately for train (blue, solid) and validation (red, dashed) splits, for the unmasked (left) and masked (right) models. A narrower train distribution relative to the validation distribution indicates over-fitting.

**Supplementary Table A1.** Performance summary of all models before and after bias correction. The results show minimal changes overall, with a general trend of improved performance after correction.

| Dataset | Before correction |  |  | After correction |  |  |
| --- | --- | --- | --- | --- | --- | --- |
|  | Train MAE | Val MAE | Test MAE | Train MAE | Val MAE | Test MAE |
| Whole body | $1.754 \pm 1.12$ | $2.639 \pm 1.21$ | $2.719 \pm 1.26$ | $1.706 \pm 1.31$ | $2.492 \pm 1.37$ | $2.564 \pm 1.45$ |
| Brain | $1.657 \pm 1.64$ | $2.644 \pm 1.34$ | $2.906 \pm 1.73$ | $1.500 \pm 1.62$ | $2.338 \pm 1.35$ | $2.496 \pm 1.75$ |
| Heart | $1.909 \pm 4.16$ | $3.572 \pm 4.11$ | $3.405 \pm 2.20$ | $0.980 \pm 1.67$ | $2.634 \pm 2.38$ | $3.022 \pm 3.34$ |
| Spine | $0.789 \pm 1.39$ | $3.164 \pm 1.70$ | $3.279 \pm 2.19$ | $0.750 \pm 1.32$ | $3.028 \pm 1.71$ | $3.164 \pm 2.44$ |
| Liver | $0.882 \pm 1.45$ | $4.014 \pm 1.83$ | $4.081 \pm 1.85$ | $0.841 \pm 1.42$ | $3.842 \pm 1.87$ | $3.929 \pm 1.87$ |
| Lungs | $0.802 \pm 1.16$ | $3.166 \pm 1.56$ | $3.255 \pm 1.60$ | $0.784 \pm 1.15$ | $3.071 \pm 1.57$ | $3.152 \pm 1.60$ |
| Muscle | $1.001 \pm 1.24$ | $2.702 \pm 1.28$ | $2.847 \pm 1.34$ | $0.939 \pm 1.20$ | $2.627 \pm 1.30$ | $2.722 \pm 1.36$ |
| Intestine | $1.056 \pm 1.52$ | $3.751 \pm 1.92$ | $3.798 \pm 2.01$ | $1.033 \pm 1.51$ | $3.639 \pm 1.93$ | $3.668 \pm 2.02$ |

### A.2 Chronic Diseases

We summarise the results of our analysis comparing accelerated ageing and specific diseases for all body regions in Figure A4. We note that corresponding visualisations for the whole-body age and the brain age are contained in the main part of this manuscript.

### A.3 Lifestyle Features

Figure A5 shows the differences in predicted age versus chronological age for smokers (red) and non-smokers (blue) for different body region ages (heart, liver, spine, muscle, intestine). Smokers tend to show more accelerated ageing compared to non-smokers. Their age prediction lies mostly above the line of perfect prediction (orange), while non-smokers are more evenly distributed showing accelerated and decelerated ageing.

### A.4 Survival Analysis

We visualise additional Kaplan Meier curves for the remaining body regions that show the difference in survival probability between accelerated agers (“acc”, orange) and decelerated agers (“dec”, blue) in Figure A6. The shaded area indicates the confidence interval. For all visualised body regions (lungs, liver, spine, muscle, intestine, heart), 5106 subjects show accelerated ageing and 5234 decelerated ageing.

### A.5 Virtual Ageing Model

We here summarise additional results of the Virtual Ageing Model, where we simulate younger body region ages for all regions individually as well as for all accelerated regions and summarise the respective whole-body ages and the corresponding age gaps in Table A2. We highlight that artificially reducing the spine age has the same impact on the whole-body age as simulating decelerated ages for all accelerated body regions. With these experiments, it always needs to be kept in mind that even though a simulated younger region does not impact the overall whole-body age prediction, this can still mean that it is important for the subject to intervene on their body-region ages and, for example, make corresponding lifestyle changes. In this experiment, replacing the spine in the virtual ageing model led to the least accelerated whole-body age with an age gap of +4.06.

**Supplementary Table A2.**
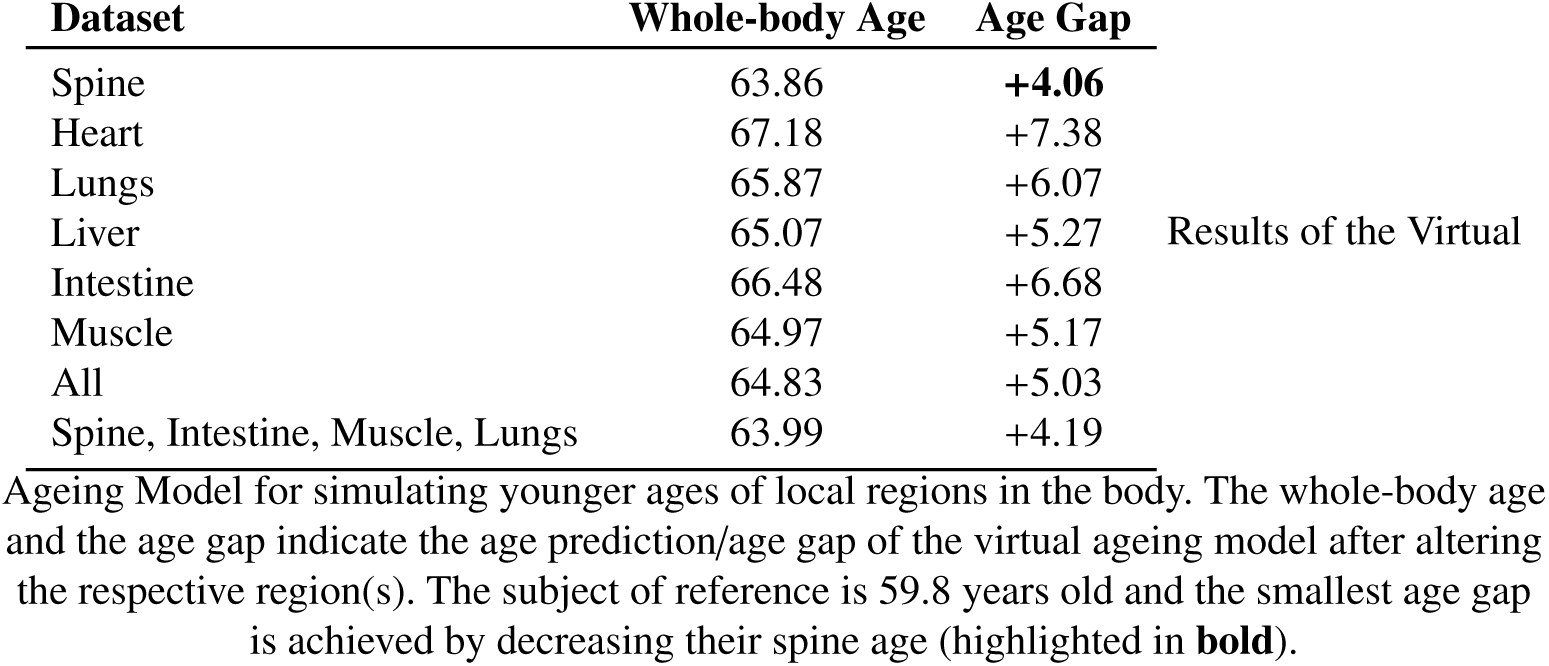
Impact of regional virtual ageing on whole-body age predictions.

**Supplementary Figure A2.**
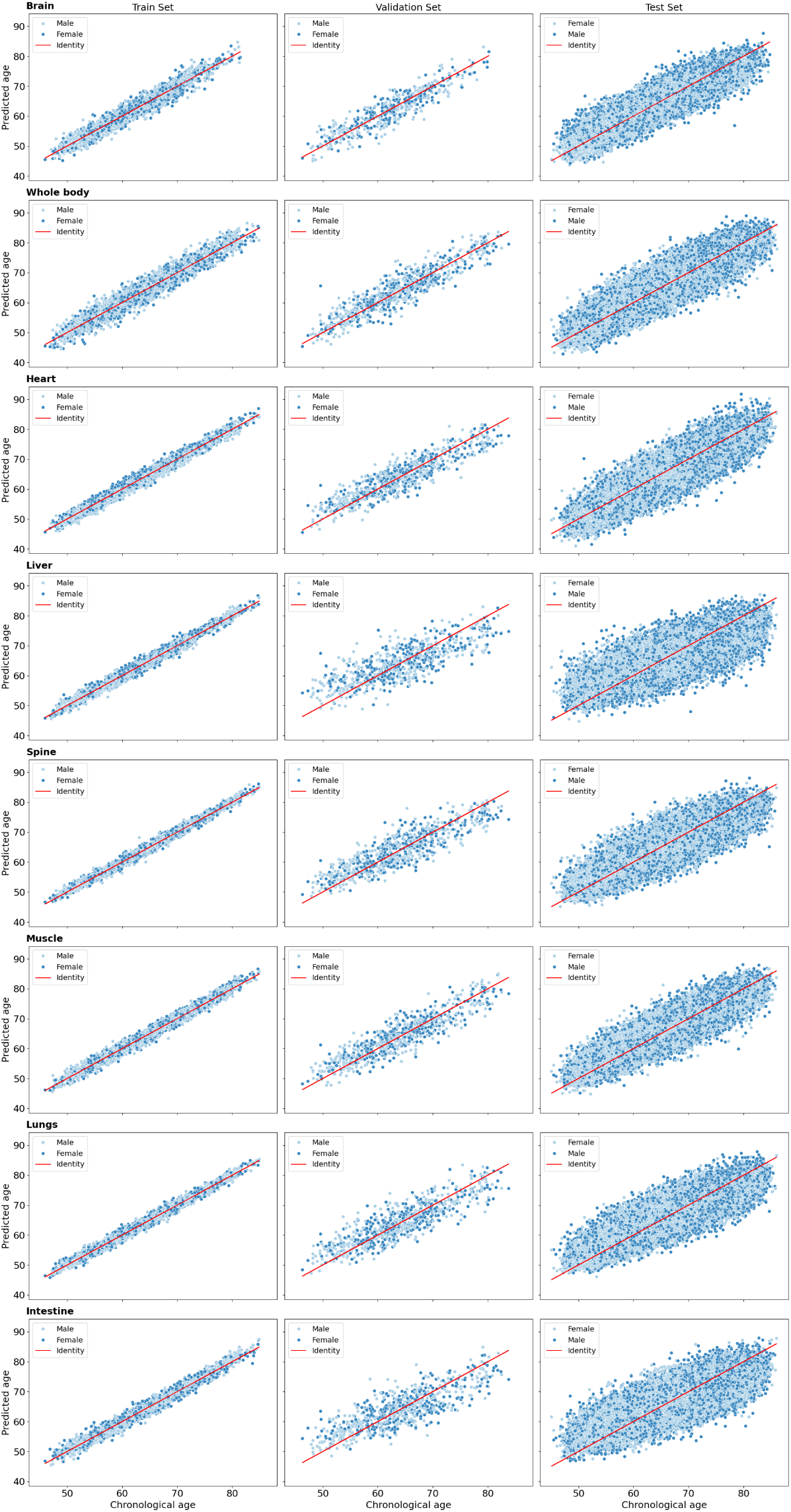
Overall model performances. Model performance on the train set (left), the validation set (middle), and the test set (right) for the brain (first row), whole body (second row), heart (third row), liver (fourth row), spine (fifth row), muscle (sixth row), lungs (seventh row), and intestine (eighth row). The red line indicates a perfect prediction and the colours indicate sex (blue: female, dark blue: male).

**Supplementary Figure A3.**
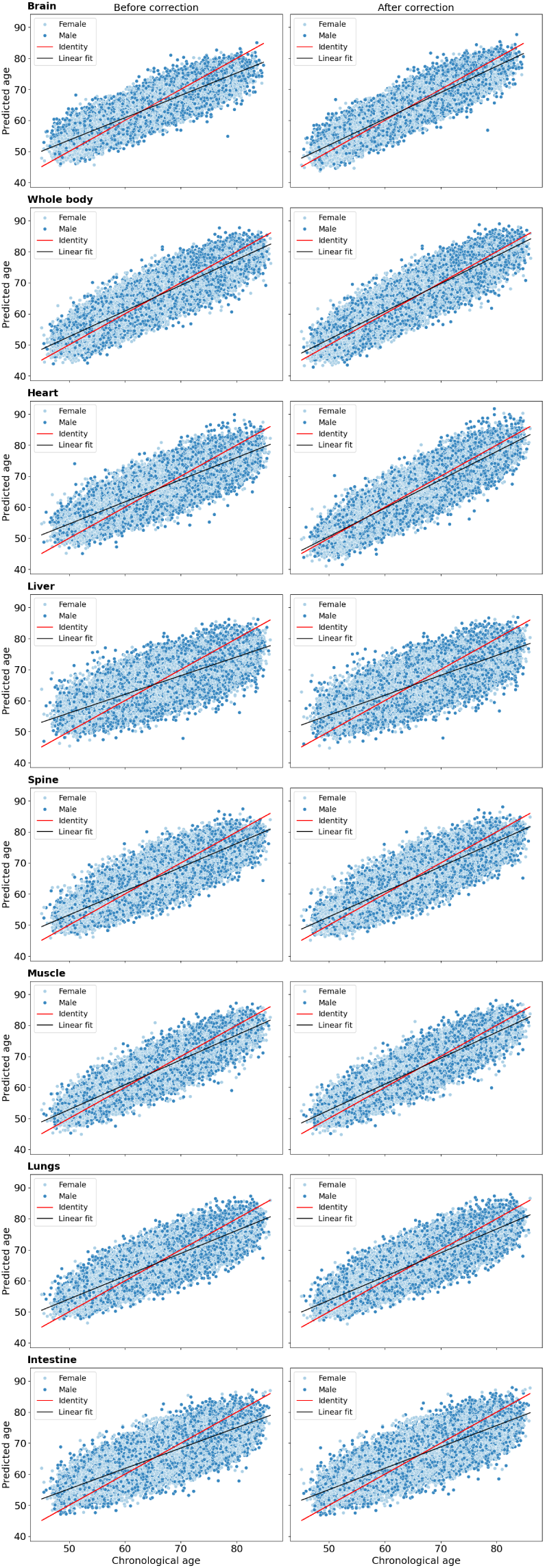
Effect of bias correction on model predictions. Predicted age vs. chronological age before bias correction (left) and after bias correction (right) for the brain (first row), whole body (second row), heart (third row), liver (fourth row), spine (fifth row), muscle (sixth row), lungs (seventh row), and intestine (eighth row). The red line indicates a perfect prediction, the black line shows a linear fit to the data, and the colours indicate sex (blue: female, dark blue: male).

**Supplementary Figure A4.**
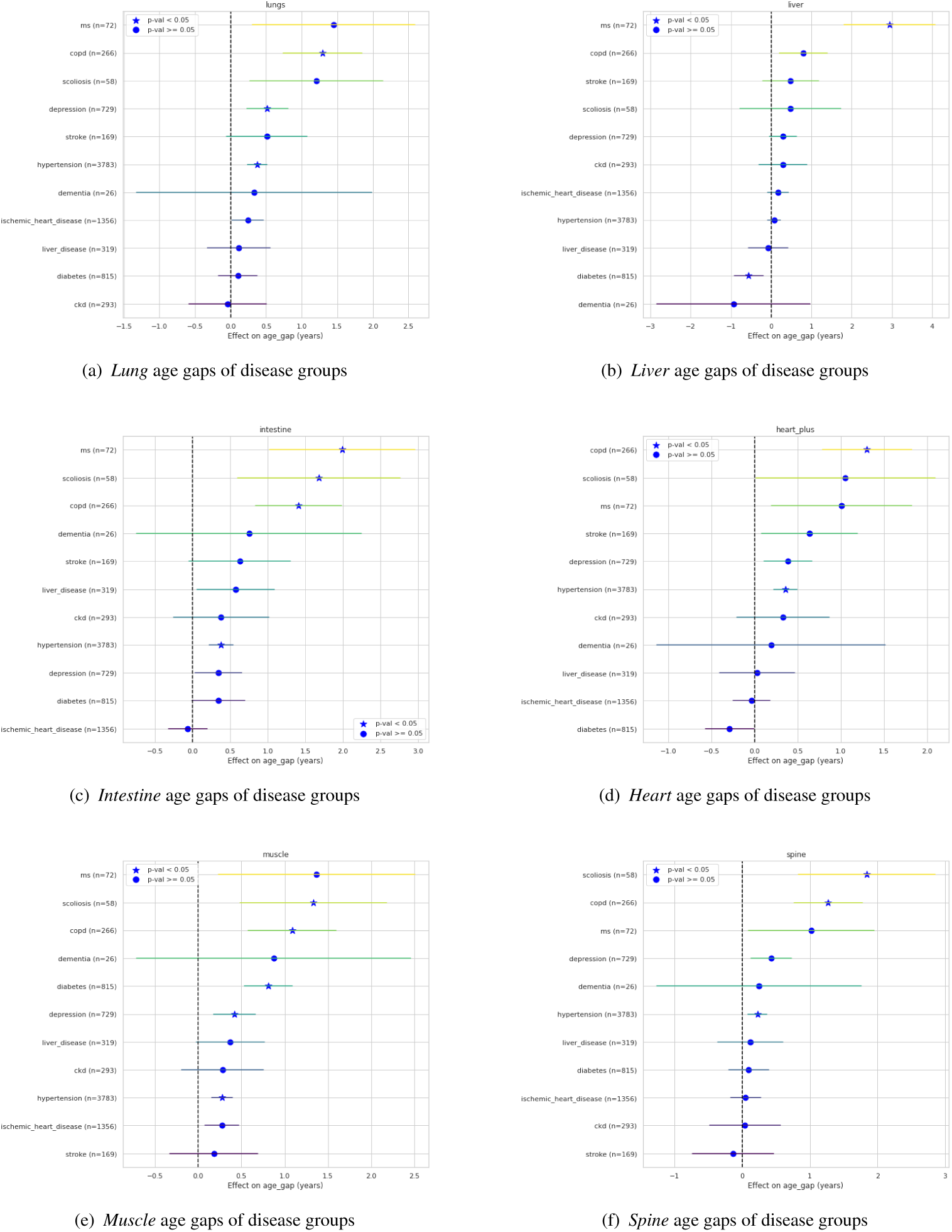
Propensity-weighted age gaps across diseases for brain and whole-body datasets. Distribution of age gaps for different diseases for (a) the lung, (b) the liver, (c) the intestine, (d) the heart, (e) the muscle, and (f) the spine age predictions. The dotted horizontal (red) line indicates the average age gap of all subjects with diseases (test set). The diseases are ordered by ascending median age gap.

**Supplementary Figure A5.**
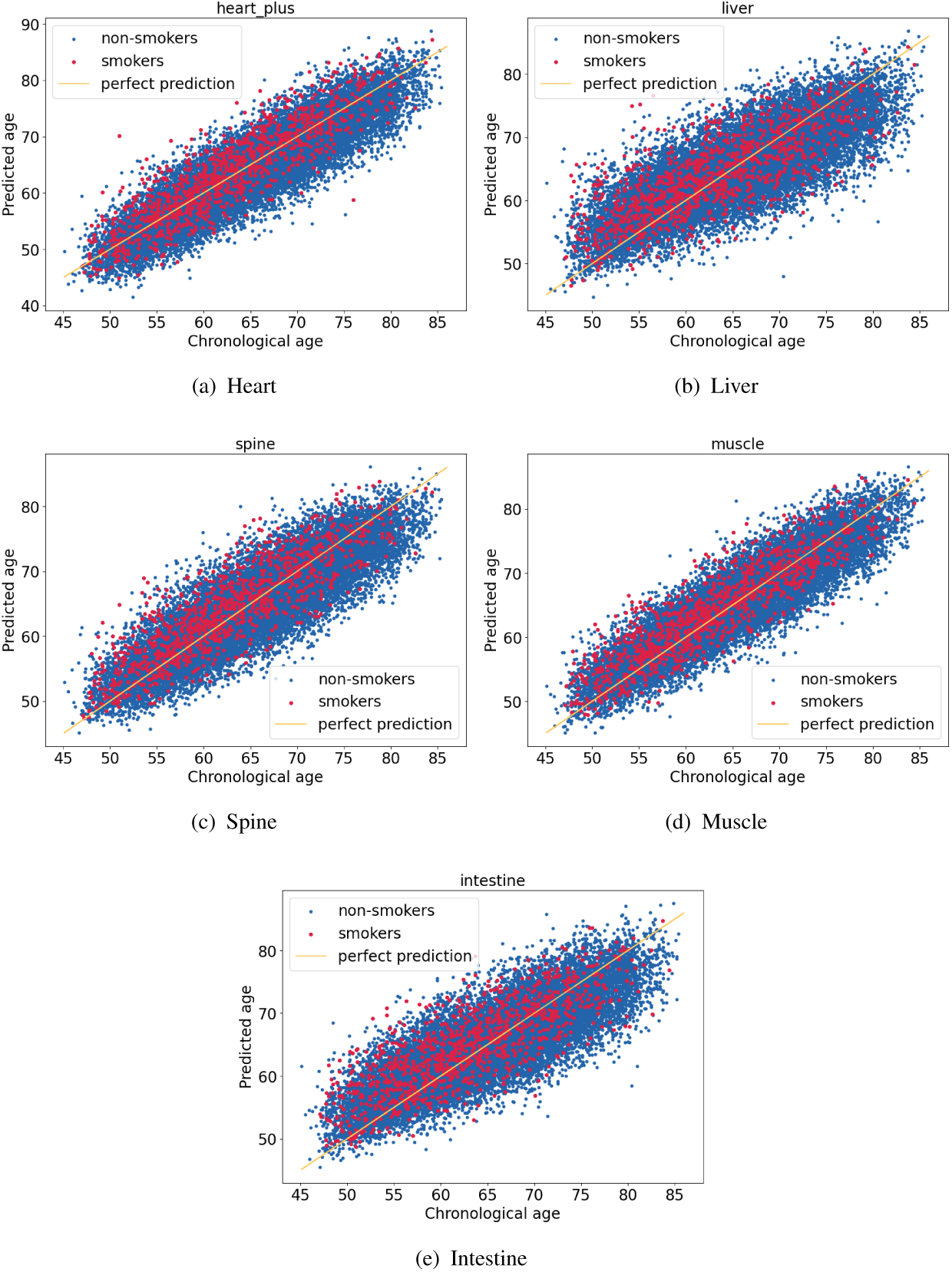
Body region age predictions by smoking status. Visualisation of the predicted age vs. the chronological age of smokers (red) and non-smokers (blue) for (a) the heart, (b) the liver, (c) the spine, (d) the muscle, and (e) the intestine images. Smokers tend to show more accelerated ageing in all image datasets.

**Supplementary Figure A6.**
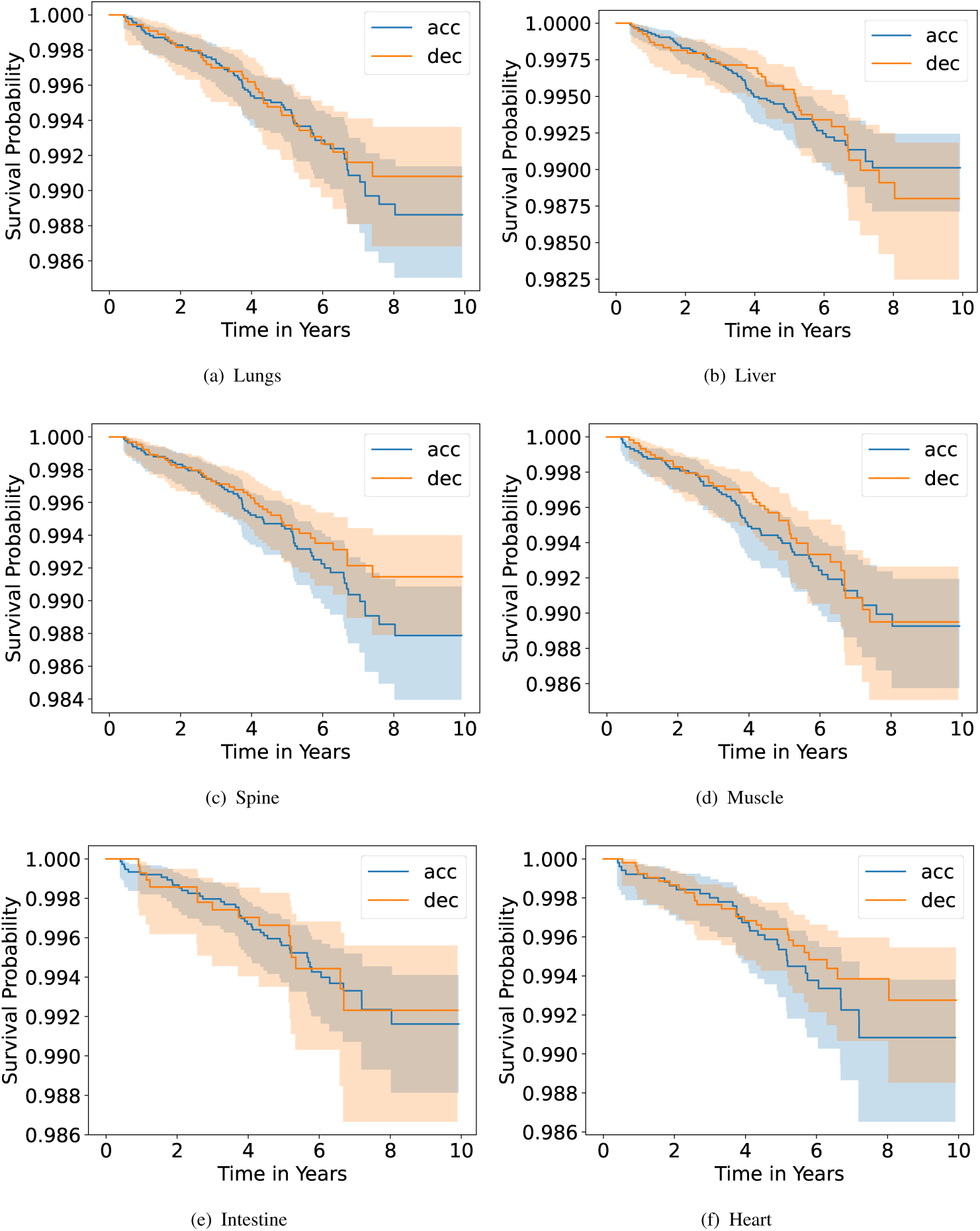
Kaplan Meier curves for body-region ages. This figure compares accelerated (acc, blue) and decelerated (dec, orange) agers for the lungs (a), the liver (b), the spine (c), the muscles (d), the intestine (e) and the heart (f) with the associated error bands.

## Supplementary Material B Dataset details

### B.1 Summary Dataset

**Supplementary Table B3.** Baseline characteristics of the whole body dataset.

| Characteristic | Females (n=26,331) | Males (n=23,335) |
| --- | --- | --- |
| Age (years) | 64.8 ± 5.2 | 65.5 ± 5.2 |
| Weight (kg) | 69.3 ± 13.4 | 84.2 ± 13.6 |
| Height (cm) | 163.5 ± 6.4 | 177.2 ± 6.7 |
| Diabetes | 2.5% | 4.4% |
| Stroke | 0.5% | 1.0% |
| Multiple Sclerosis (MS) | 0.4% | 0.1% |
| Dementia | 0.1% | 0.1% |
| COPD | 1.1% | 1.2% |
| Chronic Kidney Disease (CKD) | 1.2% | 1.3% |
| Hypertension | 13.0% | 19.6% |
| Ischemic Heart Disease | 3.1% | 8.5% |
| Depression | 3.8% | 2.3% |
| Scoliosis | 0.3% | 0.1% |
| Liver Disease | 1.3% | 1.6% |

**Supplementary Table B4.** Baseline characteristics of the brain dataset.

| Characteristic | Females (n=17,350) | Males (n=15,128) |
| --- | --- | --- |
| Age (years) | 64.7 ± 5.2 | 65.1 ± 5.2 |
| Weight (kg) | 68.9 ± 13.0 | 83.4 ± 13.3 |
| Height (cm) | 163.5 ± 6.4 | 176.9 ± 6.7 |
| Diabetes | 2.4% | 3.9% |
| Stroke | 0.5% | 0.8% |
| Multiple Sclerosis (MS) | 0.3% | 0.1% |
| Dementia | 0.1% | 0.1% |
| COPD | 1.1% | 1.1% |
| Chronic Kidney Disease (CKD) | 1.1% | 1.1% |
| Hypertension | 12.4% | 18.2% |
| Ischemic Heart Disease | 3.1% | 7.8% |
| Depression | 3.2% | 2.0% |
| Scoliosis | 0.3% | 0.1% |
| Liver Disease | 1.3% | 1.5% |

### B.2 Dataset overlap

Figure B7 describes the overlap, in terms of subjects, of all datasets (train, val and test).

**Supplementary Figure B7.**
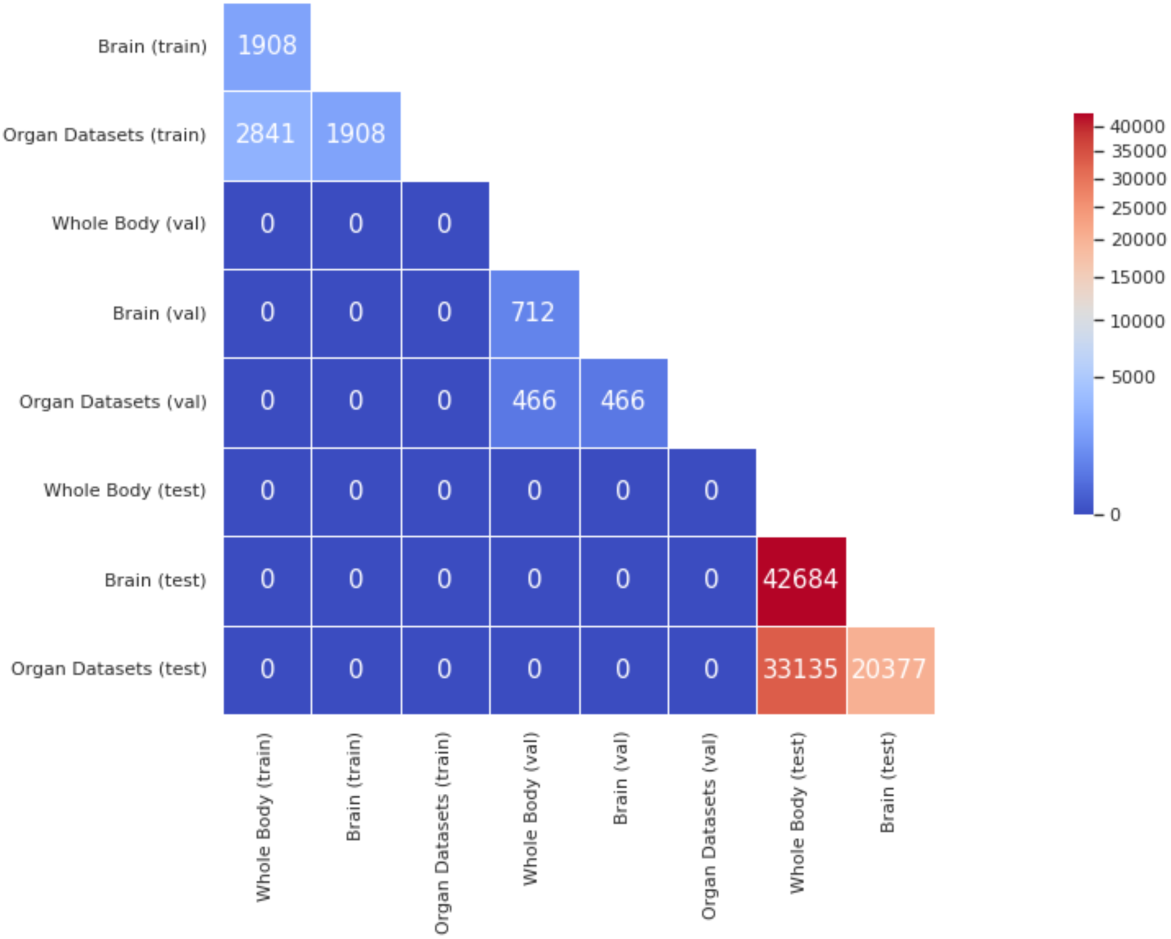
Subject overlap of all datasets. Brain subjects are included in the whole-body and organ datasets. Overlap exists within training, validation, and test sets, but not across splits.

### B.3 Age gap distribution

Figure B8 describes the predicted age gap distribution by sex for all datasets.

**Supplementary Figure B8.**
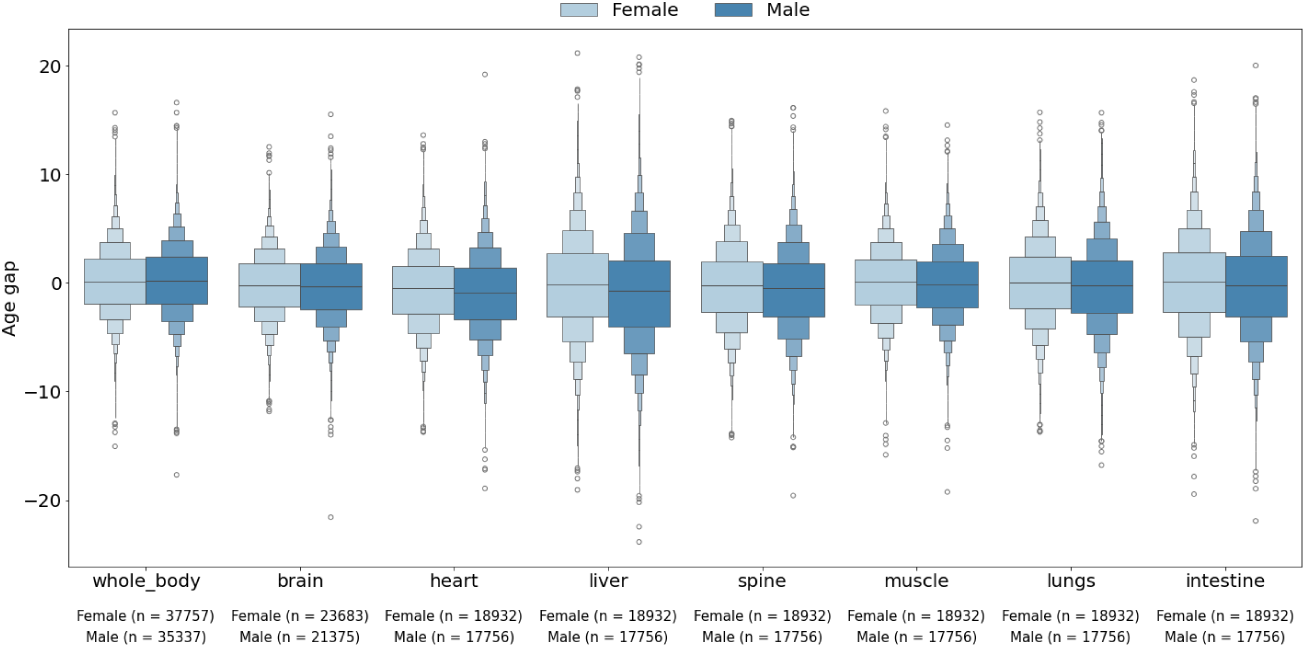
Age gap distributions for all body region images for male and female subjects.

